# Exposure to environmental level pesticides stimulates and diversifies evolution in *Escherichia coli* towards greater antibiotic resistance

**DOI:** 10.1101/665273

**Authors:** Yue Xing, Shuaiqi Wu, Yujie Men

## Abstract

Antibiotic resistance is one of the most challenging issues in public health. Antibiotic resistance can be selected by antibiotics at sub-inhibitory concentrations, the concentrations typically occurring in natural and engineered environments. Meanwhile, many other emerging organic contaminants such as pesticides are frequently co-occurring with antibiotics in agriculture-related environments and municipal wastewater treatment plants. To investigate the effects of the co-existing, non-antibiotic pesticides on the development of antibiotic resistance, we conducted long-term exposure experiments using a model *Escherichia coli* strain. The results revealed that 1) the exposure to a high level (in mg/L) of pesticides alone led to the emergence of mutants with significantly higher resistance to streptomycin; 2) the exposure to an environmental level (in µg/L) of pesticides together with a sub-inhibitory level (in sub mg/L) of ampicillin synergistically stimulated the selection of ampicillin resistance and the cross-selection of resistance to three other antibiotics (i.e., ciprofloxacin, chloramphenicol, and tetracycline). Resistance levels of mutants selected from co-exposure were significantly higher than those of mutants selected from ampicillin exposure only. The comparative genomic and transcriptomic analyses indicate that distinct and diversified genetic mutations in ampicillin- and ciprofloxacin-resistant mutants were selected from co-exposure, which likely caused holistic transcriptional regulation and the increased antibiotic resistance. Together, the findings provide valuable fundamental insights into the development of antibiotic resistance under environmentally relevant conditions, as well as the underlying molecular mechanisms of the elevated antibiotic resistance induced by the exposure to pesticides.

**Significance statement:** Antibiotic resistance is a major threat to public health globally. Besides clinically relevant environments, the emergence and spread of resistant bacteria in non-clinical environments can also potentially pose risks of therapy failures. This study showed that the long-term, environment-level exposure to pesticides with and without antibiotics significantly stimulated the development of greater antibiotic resistance. The resistant strains selected from the exposure to pesticides are genetically and metabolically distinct from the ones selected by the antibiotic only. Although it is still being debated regarding whether or not a large use of antibiotics in plant agriculture is harmful, our findings provide the first fundamental evidence that greater concerns of antibiotic resistance may result if antibiotics are applied together with non-antibiotic pesticides.

## Introduction

Antibiotic resistance has been one of the most challenging environmental and public health issues. The *de novo* mutation is one important route for bacteria to acquire antibiotic resistance, under both clinical and environmental conditions (1-4). Antibiotics at both inhibitory (typically in the mg/L range) and sub-inhibitory concentrations (below the minimal inhibitory concentration (MIC); in the high µg/L or ng/L range) can lead to increased resistance emergence (1, 3, 5). The latter is of more concern due to the ubiquitous occurrence of antibiotic residues at low (i.e., sub-MIC) levels in the environment. This may explain the emergence of antibiotic resistance in many non-clinically relevant environments such as domestic sewage, water bodies receiving treated sewage from municipal wastewater treatment plants (WWTPs), as well as farm run-off where antibiotics occur from tens of ng/L to several hundreds of μg/L (6-9). Although there is limited direct evidence in terms of resistant phenotypes, the wide surveillance of antibiotic resistance genes (ARG) revealed elevated ARG levels in treated wastewater and the receiving environment along with the occurrence of low-level antibiotics (10-13). In addition, cross-selection occurs in mutants exposed to a single antibiotic at sub-MIC levels, which developed resistance not only to the exposed antibiotic but also to other non-exposed antibiotics, indicating the complexity of antibiotic resistance development (14, 15). With similar cytotoxicity to antibiotics, disinfectants and disinfection byproducts were found to promote the development of antibiotic resistance at high levels (several to a few thousand mg/L) (16-19).

Moreover, in natural and built environments a variety of other emerging organic contaminants such as pesticides, non-antibiotic drugs, and personal care products are usually co-occurring with antibiotics at low levels (20-24). It is still unclear how these non-antibiotic emerging organic contaminants at environmentally relevant concentrations would affect the selection of antibiotic resistance by antibiotics. If synergistic effects were present, antibiotic resistance levels in those environments would be underestimated by only considering the antibiotic occurrence. Thus, it is crucial to obtain a better understanding of the emergence of antibiotic resistance with exposure to both antibiotics and non-antibiotic organic contaminants at environmental levels.

Pesticides are one important group among those contaminants co-existing with antibiotics. They are typically found in agricultural soils, run-offs, and the receiving water bodies (25-28), which are also potentially antibiotic-impacted environments. For example, antibiotics used in farms to treat sick animals and boost livestock growth can be released into those environments (8, 29), leading to the co-occurrence of pesticides and antibiotics. Moreover, antibiotics have also been applied to fight against plant bacterial diseases in plant farms (30), where pesticides may also be used. For instance, currently the US Environmental Protection Agency is in the process of allowing heavy usage of two antibiotics, streptomycin and oxytetracycline, to combat citrus greening, a bacterial disease killing citrus trees (31). In addition, pesticides also occurred in other non-clinical antibiotic-impacted environments such as irrigation water and municipal wastewater (32-37). The environmental levels of each individual pesticide range from less than 1 ng/L to tens of μg/L (25, 26), which brings an overall environmental occurrence to a high µg/L range due to the presence of many pesticide species used for different purposes in farms and households. Some pesticides like biocides share similar inhibitory mechanisms with antibiotics, such as membrane disruption (38-40) and inhibition of cell wall synthesis (41, 42), which might favor mutations towards the co-selection of antibiotic resistance.

The goal of this study was to fill the knowledge gap regarding the emergence of antibiotic resistance in environments with the occurrence of both antibiotics and pesticides. We aimed to 1) investigate the effects of environmental level exposure to pesticides alone and the interactive effects with sub-MIC antibiotics (synergistic, antagonistic, or neutral) on the development of antibiotic resistance, and 2) identify the underlying mechanisms of the emerged antibiotic resistance from 1). To accomplish our goals, we designed evolutionary experiments with a susceptible *Escherichia coli* strain being exposed to pesticides only and co-exposed to ampicillin and pesticides at environmental levels for 500 generations. The change in antibiotic resistance of *de no* mutants was determined. Genetic mutations of the *de no* mutants selected from different exposure conditions were identified by whole-genome sequencing (WGS). Transcriptional regulation in resistant mutants from co-exposure compared to those from antibiotic exposure alone was examined using RNA sequencing (RNA-seq) and reverse transcription quantitative PCR (RT-qPCR). The genomic and transcriptomic analyses provide insights into different mechanisms of antibiotic resistance developed under those exposure conditions.

## Results

### Effects of the Exposure to Pesticides on the Development of Streptomycin Resistance

We initiated evolutionary experiments with a susceptible strain *E. coli* K12 C3000. Starting from one ancestor strain (G0), parallel population passages were exposed to constant concentrations of pesticide mixture. The mixture consisted of 23 frequently detected pesticides in various natural and engineered environments, including biocides, fungicides, herbicides, and insecticides (Table S1). The concentrations of pesticides used in this study were based on their environmental concentrations (EC) (0.1 – 4.8 µg/L, each; ∼ 20 µg/L in total) (Table S1). The total pesticide concentrations used for the exposure experiments range from 1/125EC (0.16 µg/L) to 125EC (2.5 mg/L). In the control experiments, *E. coli* passages were obtained from the same ancestor cells in the same way but without pesticide exposure.

Among all pesticide exposure levels, an increased mutation frequency (1 – 2 orders of magnitude higher) towards streptomycin resistance was observed in *E. coli* populations being exposed to 125EC pesticides for 500 generations (G500) (Figure S1), although with no statistical significance due to the large variations among triplicated evolution passages. This indicates that exposure to high-level (2.5 mg/L) pesticides stimulated the emergence of streptomycin (Strep) resistance. To further characterize the resistance levels of Strep-resistant mutants, 36 resistant mutants were randomly picked up on selective solid media, and their MICs were determined. Mutants selected from 125EC exposure acquired significantly higher Strep-MICs compared to the mutants from the no-exposure control (*p*-value = 0.0013, N = 36, Mann-Whitney U test) (Figure 1). It is worth noting that the exposure to high-level pesticides favored the emergence of *de novo* mutants more resistant to Strep only, but not to the other four tested antibiotics, i.e., ampicillin (Amp), tetracycline (Tet), ciprofloxacin (Cip), and chloramphenicol (Chl).

**Fig. 1.**
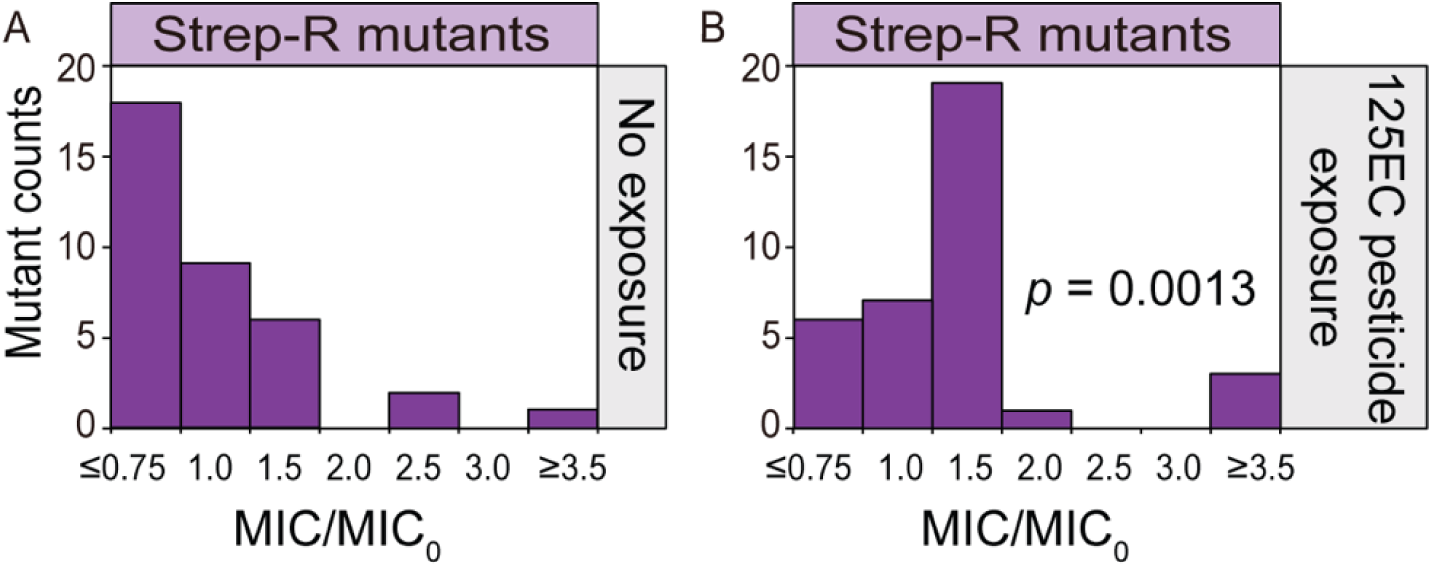
MICs of non-exposed mutants from G500 (A) and mutants from high-level pesticide exposure (B). MIC_0, Strep_ is the MIC (9 mg/L) of the ancestor strain. The *p* value of Mann-Whitney U test is indicated (n = 36).

### Effects of Pesticides + Ampicillin Co-exposure at Environmental Levels on the Development of Antibiotic Resistance

We next tested the combined effects of environmental level pesticides and Amp (1/125 – 1/5 MIC_0_) on the development of antibiotic resistance in *E. coli*. According to the MIC distribution of the 36 Amp-resistant mutants, those selected from populations co-exposed to 1/5MIC_0_ Amp and 1EC pesticides exhibited a shift to higher MICs (Figure 2B), compared to those selected from populations exposed to 1/5MIC_0_ Amp (Figure 2A). The shift of MIC distribution was statistically significant (*p*-value = 0.039, Mann-Whitney U test). To explore the development of cross-resistance, we determined the MIC distributions of *E. coli* mutants from co-exposed and Amp-exposed populations (G500) resistant to four other antibiotics: Strep, Chl, Cip, and Tet. Except for Strep with similar resistance developed under co- and Amp-exposure conditions, the resistance (i.e., MICs) to the other three antibiotics were significantly higher (1.5 – 3.5 times) for resistant mutants from co-exposure than from Amp-exposure (*p*-values of 1.1 × 10^-10^, 0.044, and 1.3 × 10^-7^ for Chl, Cip, and Tet, respectively) (Figure 2 D, F and H). The co-exposure also accelerated the development of resistance to the four antibiotics with mutation frequencies 1 – 4 orders of magnitude higher than those under Amp-exposure only, although without statistical significance due to variations among the triplicated evolution passages (Figure S2). Collectively, the co-exposure to environmental level pesticides and 1/5MIC_0_ ampicillin exhibited synergistic effects on the emergence of mutants resistant to not only the exposed antibiotic but also other non-exposed antibiotics (cross-selection). The co-exposure condition selected mutants more resistant than those selected under sub-MIC antibiotic selection pressure only. One should note that no accelerated development of higher resistance was observed for G500 *E. coli* cells exposed to the same concentration of pesticides but with lower ampicillin concentrations (1/125MIC_0_ and 1/25MIC_0_), as well as the G500 non-exposure control (data not shown).

**Fig. 2.**
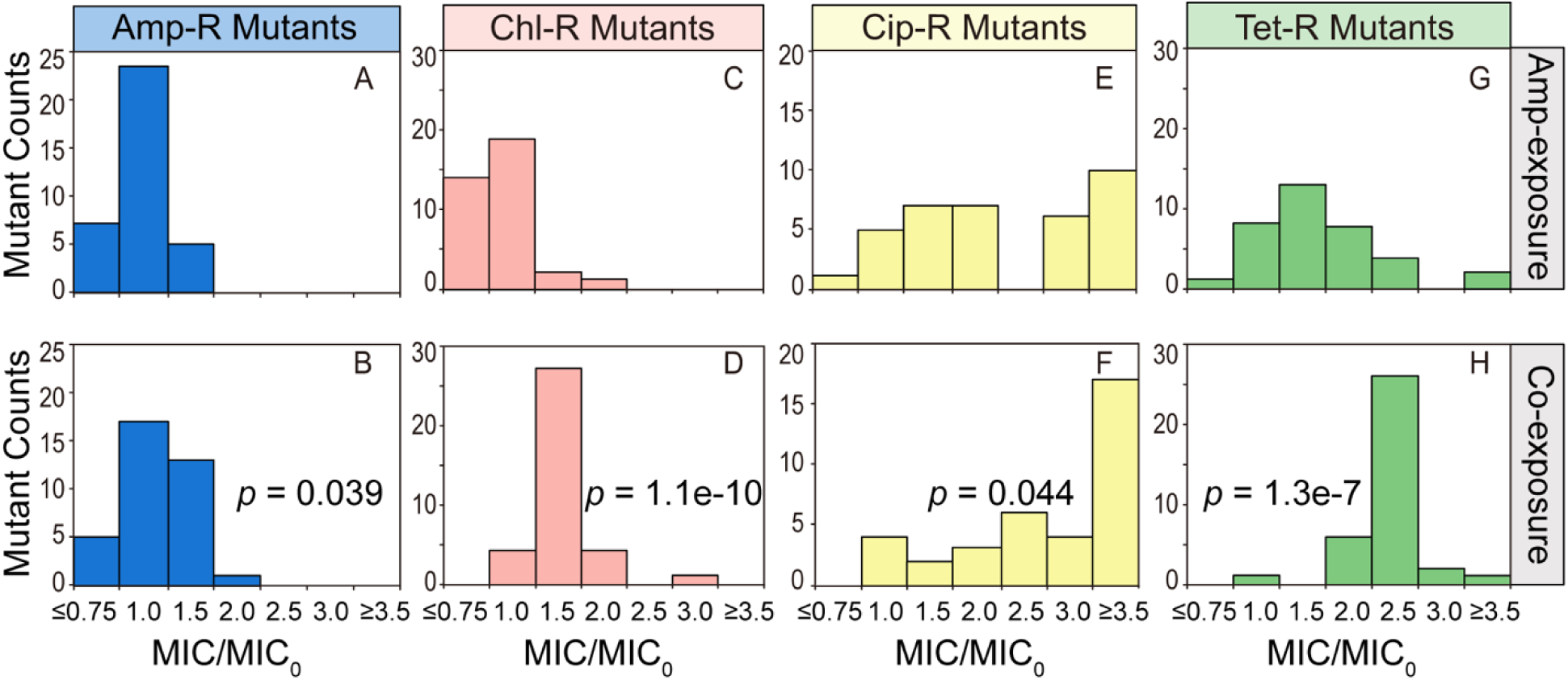
MICs of resistant mutants selected from G500 *E. coli* under co-exposure (1EC pesticides + 1/5MIC_0, Amp_) (bottom-row; A: Amp-R mutants, C: Chl-R mutants, E: Cip-R mutants, and G: Tet-R mutants) and single-exposure (1/5MIC_0, Amp_) (top-row; B: Amp-R mutants, D: Chl-R mutants, F: Cip-R mutants, and H: Tet-R mutants). MIC_0, Amp_ = 4 mg/L, MIC_0, Chl_ = 8 mg/L, MIC_0, Cip_ = 0.016 mg/L, and MIC_0, Tet_ = 1 mg/L. The *p* value of the Mann-Whitney U test between Amp-exposure and co-exposure conditions is indicated (n = 36).

### Genetic Mutations in Strep-resistant Mutants Selected by High-level Pesticide Exposure

To unravel the mechanisms leading to the elevated mutation frequency towards Strep resistance and the higher Strep resistance of *E. coli* mutants after being exposed to high-level pesticides (125EC), we identified valid genetic mutations including non-synonymous single nucleotide polymorphisms (SNPs), insertions and deletions (INDEL) in the resistant mutants compared to the susceptible strains from G500 population without pesticide exposure, which have the same MIC as the ancestor strain (G0).

The genomes of Strep-resistant mutants isolated from G500 *E. coli* with pesticide exposure revealed genetic mutations in four genes including two SNPs, and two deletions against the genome of the susceptible isolate from G500 *E. coli* without pesticide exposure (Table 1, see Table S3 in the SI for a complete list of genetic mutations). These four mutated genes encode proteins for: (i) target modification; (ii) DNA replication and repair; (iii) regulation. It is noteworthy that all three sequenced Strep-resistant mutants from the pesticide-exposed cultures shared the same genetic mutation of the *rpsG* gene (SNP: A → T), resulting in gaining a stop codon (*) replacing Leu157 in the amino acid sequence (Table 1). The *rpsG* gene encodes the 30S ribosomal protein S7, which is essential for cell growth. The stop codon gained at a later position (157 of 179 residues) of the amino acid sequence did not affect the function of this protein, as no growth inhibition of the resistant mutants was observed (data not shown).

**Table 1.**
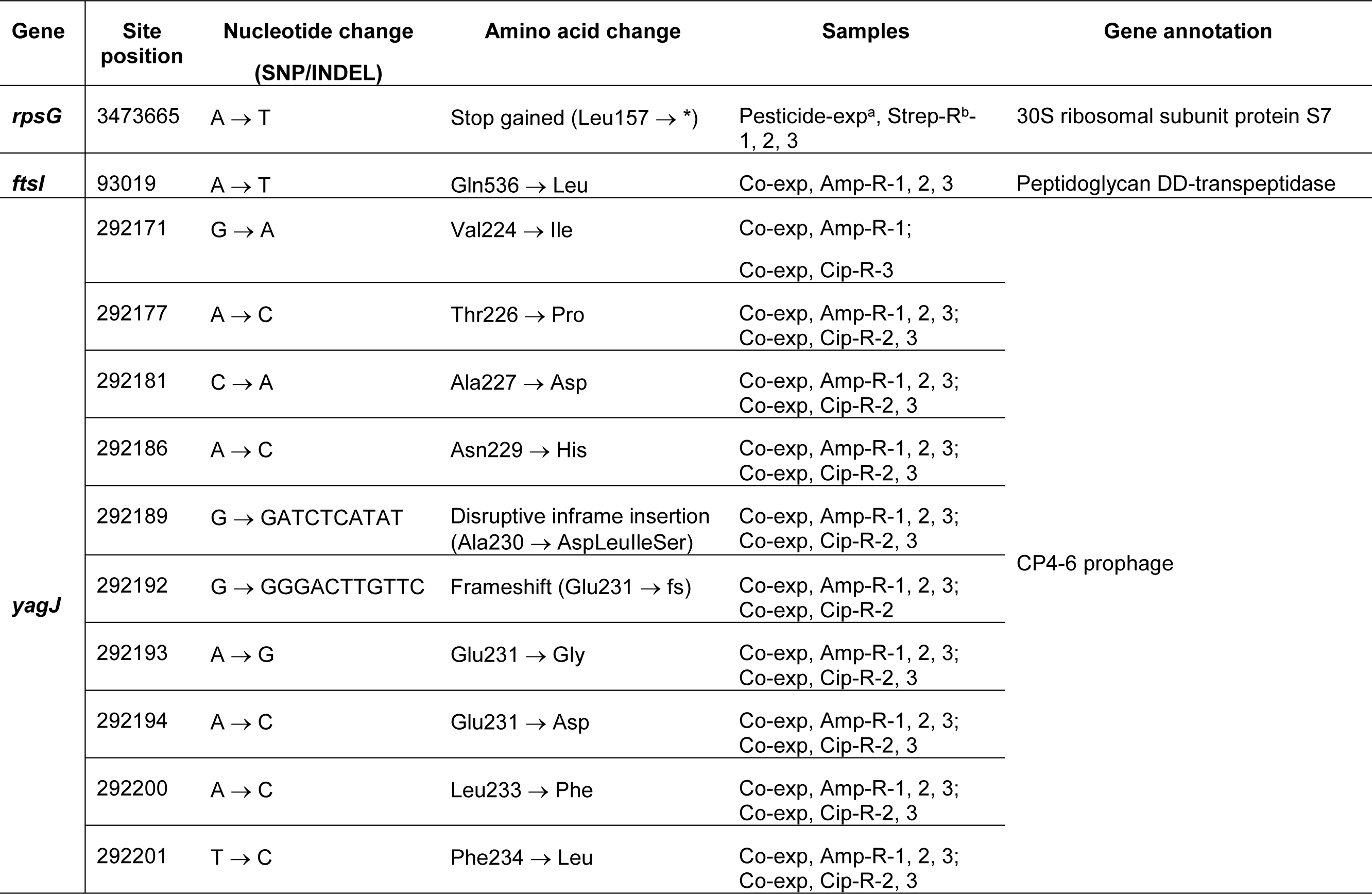
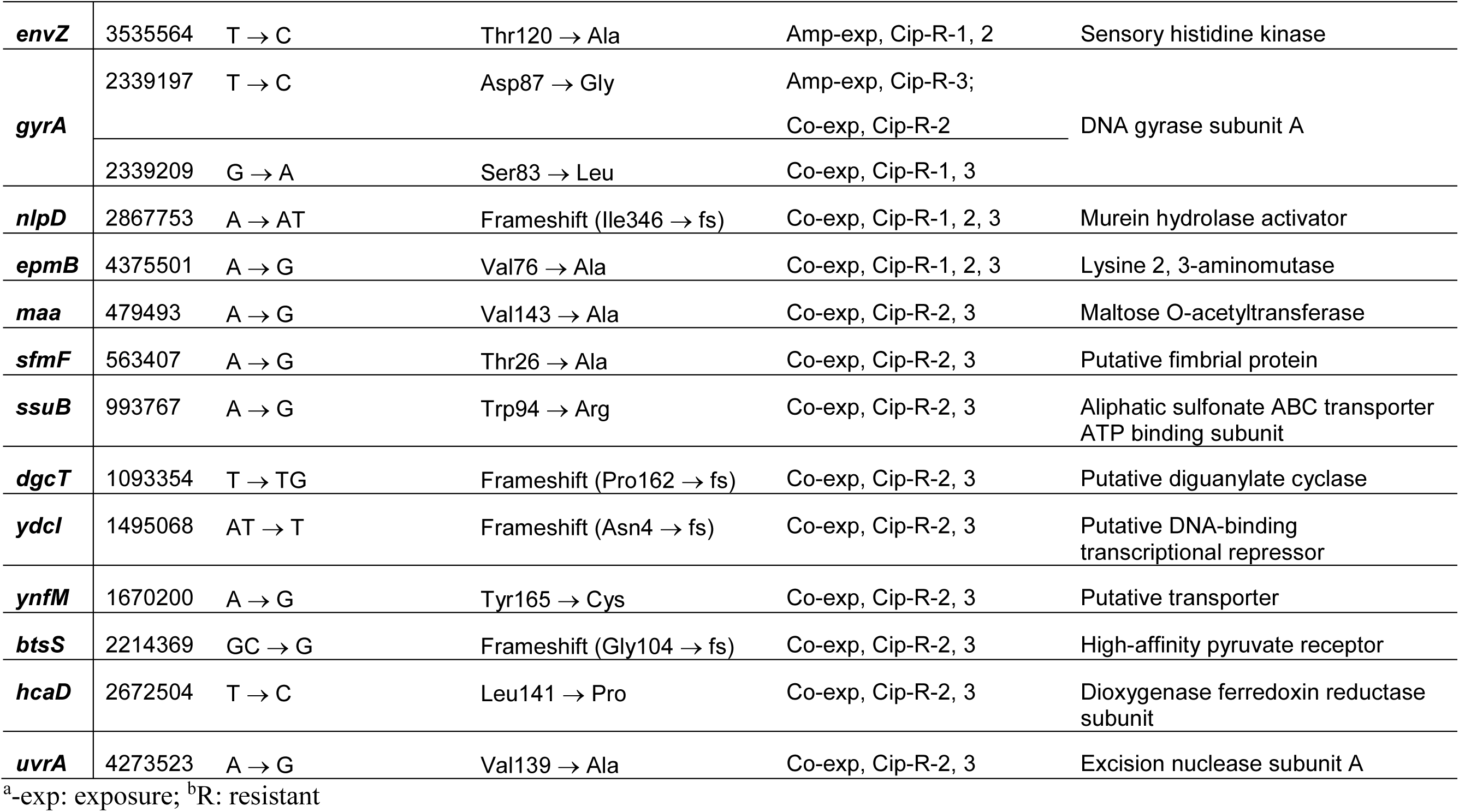
Selected genetic mutations identified in the resistant mutants.

| Gene | Site position | Nucleotide change<br>(SNP/INDEL) | Amino acid change | Samples | Gene annotation |
| --- | --- | --- | --- | --- | --- |
| *rpsG* | 3473665 | A → T | Stop gained (Leu157 → *) | Pesticide-exp<sup>a</sup>, Strep-R<sup>b</sup>-1, 2, 3 | 30S ribosomal subunit protein S7 |
| *ftsI* | 93019 | A → T | Gln536 → Leu | Co-exp, Amp-R-1, 2, 3 | Peptidoglycan DD-transpeptidase |
| *yagJ* | 292171 | G → A | Val224 → Ile | Co-exp, Amp-R-1;<br>Co-exp, Cip-R-3 | CP4-6 prophage |
|  | 292177 | A → C | Thr226 → Pro | Co-exp, Amp-R-1, 2, 3;<br>Co-exp, Cip-R-2, 3 |  |
|  | 292181 | C → A | Ala227 → Asp | Co-exp, Amp-R-1, 2, 3;<br>Co-exp, Cip-R-2, 3 |  |
|  | 292186 | A → C | Asn229 → His | Co-exp, Amp-R-1, 2, 3;<br>Co-exp, Cip-R-2, 3 |  |
|  | 292189 | G → GATCTCATAT | Disruptive inframe insertion<br>(Ala230 → AspLeulleSer) | Co-exp, Amp-R-1, 2, 3;<br>Co-exp, Cip-R-2, 3 |  |
|  | 292192 | G → GGGACTTG TTC | Frameshift (Glu231 → fs) | Co-exp, Amp-R-1, 2, 3;<br>Co-exp, Cip-R-2 |  |
|  | 292193 | A → G | Glu231 → Gly | Co-exp, Amp-R-1, 2, 3;<br>Co-exp, Cip-R-2, 3 |  |
|  | 292194 | A → C | Glu231 → Asp | Co-exp, Amp-R-1, 2, 3;<br>Co-exp, Cip-R-2, 3 |  |
|  | 292200 | A → C | Leu233 → Phe | Co-exp, Amp-R-1, 2, 3;<br>Co-exp, Cip-R-2, 3 |  |
|  | 292201 | T → C | Phe234 → Leu | Co-exp, Amp-R-1, 2, 3;<br>Co-exp, Cip-R-2, 3 |  |
| **envZ** | 3535564 | T → C | Thr120 → Ala | Amp-exp, Cip-R-1, 2 | Sensory histidine kinase |
| **gyrA** | 2339197 | T → C | Asp87 → Gly | Amp-exp, Cip-R-3;<br>Co-exp, Cip-R-2 | DNA gyrase subunit A |
|  | 2339209 | G → A | Ser83 → Leu | Co-exp, Cip-R-1, 3 |  |
| **nlpD** | 2867753 | A → AT | Frameshift (Ile346 → fs) | Co-exp, Cip-R-1, 2, 3 | Murein hydrolase activator |
| **epmB** | 4375501 | A → G | Val76 → Ala | Co-exp, Cip-R-1, 2, 3 | Lysine 2, 3-aminomutase |
| **maa** | 479493 | A → G | Val143 → Ala | Co-exp, Cip-R-2, 3 | Maltose O-acetyltransferase |
| **sfmF** | 563407 | A → G | Thr26 → Ala | Co-exp, Cip-R-2, 3 | Putative fimbrial protein |
| **ssuB** | 993767 | A → G | Trp94 → Arg | Co-exp, Cip-R-2, 3 | Aliphatic sulfonate ABC transporter<br>ATP binding subunit |
| **dgcT** | 1093354 | T → TG | Frameshift (Pro162 → fs) | Co-exp, Cip-R-2, 3 | Putative diguanylate cyclase |
| **ydcl** | 1495068 | AT → T | Frameshift (Asn4 → fs) | Co-exp, Cip-R-2, 3 | Putative DNA-binding<br>transcriptional repressor |
| **ynfM** | 1670200 | A → G | Tyr165 → Cys | Co-exp, Cip-R-2, 3 | Putative transporter |
| **btsS** | 2214369 | GC → G | Frameshift (Gly104 → fs) | Co-exp, Cip-R-2, 3 | High-affinity pyruvate receptor |
| **hcaD** | 2672504 | T → C | Leu141 → Pro | Co-exp, Cip-R-2, 3 | Dioxygenase ferredoxin reductase<br>subunit |
| **uvrA** | 4273523 | A → G | Val139 → Ala | Co-exp, Cip-R-2, 3 | Excision nuclease subunit A |

<sup>a</sup>-exp: exposure; <sup>b</sup>R: resistant

### Genetic Mutations in Amp- and Cip-resistant Mutants Selected by the Pesticides + Ampicillin Co-exposure

To explore mechanisms leading to the higher Amp-resistance of mutants isolated in *E. coli* populations exposed to pesticides + Amp, we identified and compared the genetic mutations in Amp-resistant mutants from *E. coli* under co-exposure and Amp-exposure. We also did the same comparative genomic analysis to study the underlying mechanisms of the increase of cross-resistance to antibiotics other than the exposed Amp. We focused on Cip-resistant mutants, as they showed the highest MIC increase after the co-exposure.

For the three Amp-resistant mutants isolated from the co-exposed culture, the same mutation occurred in gene *ftsI* (SNP: A → T; amino acid change: Gln536 → Leu) (Figure 3 & Table 1). It encodes an Amp-binding protein, and this genetic mutation likely altered the protein structure, hence lowering the binding affinity of Amp to this protein. In addition, multiple mutations (non-synonymous SNPs and insertions) occurred in a prophage-related gene *yagJ*. Besides, mutations also occurred in genes encoding membrane and flagellar structure proteins (Table 1). The structural alteration of these proteins could potentially limit or avoid the entry of the antibiotic into the cells (41, 42), thus resulting in antibiotic resistance.

**Fig. 3.**
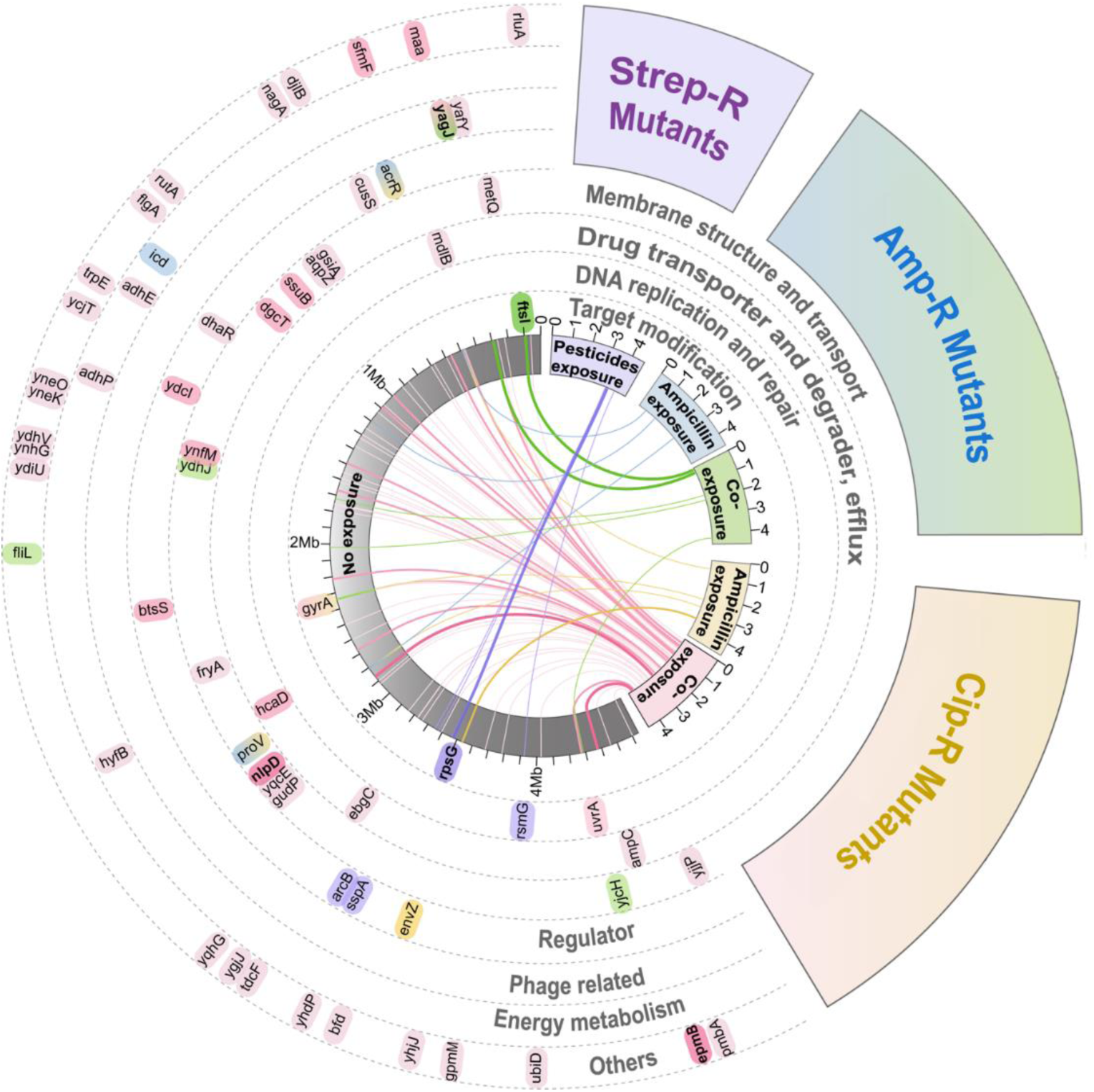
Genetic mutations identified in resistant mutants (grey: reference genome, i.e., the genome of G500 *E. coli* with no exposure; purple: genome of Strep-resistant mutants from high-level pesticide exposure; blue: genome of Amp-resistant mutant from Amp-exposure; green: genome of Amp-resistant mutant from co-exposure; yellow: genome of Cip-resistant mutants from Amp-exposure; light pink: genome of Cip-resistant mutants from co-exposure. The links represent genetic mutations (i.e., non-synonymous SNPs, insertions, or deletions). The capsules stand for mutated genes and their positions on the genome. The darker the colors and the thicker the links, the higher frequencies of the genetic mutations detected among the three resistant mutants. Rings with dashed boarders contain the genes involved in a specific function).

Interestingly, the mutations identified in Amp-resistant mutants isolated from co-exposed *E. coli* were completely different from those isolated from Amp-exposed *E. coli*. Fewer genetic mutations were detected in the Amp-resistant mutants from Amp-exposure, none of which was shared among the three sequenced mutants. One mutant had an SNP mutation in *acrR* involved in multidrug transport (43, 44) (Figure 3 & Table S3). Another mutant had an SNP mutation in the proline transport gene *proV*, which occurred in a multi-drug-resistant *Salmonella* strain (45). The third mutant had an SNP mutation in an isocitrate dehydrogenase encoding gene *icd*, the mutation of which has been observed in *E. coli* mutants resistant to nalidixic acid (46). Together, many of the identified mutated genes in the Amp-resistant strains isolated from both co-exposure and Amp-exposure conditions have resistance-related functions, which likely led to the development of Amp-resistance. Moreover, the co-exposure selected Amp-resistant mutants with distinct genetic mutations, which likely contributed to their higher MIC levels than those selected by Amp-exposure only.

For Cip-resistant mutants isolated from co-exposed *E. coli*, mutations in the *gyrA* gene occurred in all three sequenced mutants: two had Ser83 → Leu and one had Asp87 → Gly (Table 1). The DNA gyrase encoded by *gyrA* is the target of Cip, and the mutations in *gyrA* might lead to the resistance to Cip (47). Along with *gyrA* mutations, more diverse genetic mutations were detected in Cip-resistant mutants from co-exposure than from Amp-exposure, including genes with various functions: (i) DNA replication and repair; (ii) drug transporter and degrader, and efflux pumps; (iii) membrane structure and transporter; (iv) regulator; (v) prophage; and (vi) energy metabolism. Most of these mutations were not directly involved in known Cip-resistant mechanisms. It seems that the co-exposure not only accelerated but also diversified the evolution, resulting in the selection of Cip-resistant mutants with higher resistance.

There were fewer genetic mutations detected in the Cip-resistant mutants from Amp-exposure, which occurred in genes encoding proteins for target modification, transporters, and regulators (Figure 3 & Table S3). One of the three sequenced mutants had the same *gyrA* mutation (Asp87 → Gly) as the one that occurred to the Cip-resistant mutant from co-exposure condition. The other two mutants had an SNP mutation (T → C, Thr120 → Ala) in the *envZ* gene that encodes a membrane-associated protein kinase in the two-component regulatory system, which might reduce the production of membrane porin and lead to antibiotic resistance (48). The same genetic mutations of *proV* and *acrR* genes as those in the Amp-resistant mutants were found in Cip-R mutants from Amp-exposure, suggesting a more general resistance mechanism not only to ampicillin but also to other types of antibiotics.

As it is not financially applicable and practically feasible to sequence all antibiotic-resistant mutants isolated from Amp- and co-exposure for genomic comparison, complementary SNP genotyping assays were conducted to examine the prevalence of the identified genetic mutations from three biological replicates by WGS in the entire resistant population of G500 *E. coli* under co- and Amp-exposure conditions. As a representative, the *ftsI* gene, which showed the same SNP mutation among all three sequenced mutants from the co-exposure condition was targeted by the SNP genotyping assay. We treated the co-exposed and Amp-exposed G500 *E. coli* with 4 mg/L ampicillin (i.e., MIC_0, Amp_) to select resistant *E. coli* populations in the liquid media and then detected the genotyping patterns in the resistant populations. The mutated *ftsI* genotype was only detected in the resistant populations selected from co-exposed G500 *E. coli* (Figure S3). Despite the varied fractions of *ftsI* mutants in the three biological replicates (1.2%, 30.5%, and 99.8%), the presence/absence of mutated *ftsI* determined by SNP genotyping assay is consistent with the WGS results. Thus, the detection and frequency of genetic mutations from three selected mutant genomes can qualitatively represent the presence and dominance of the genotypes in the resistant population. In line with the SNP genotyping results, the replicate from co-exposure containing 99.8% mutated *ftsI* showed more than one order of magnitude higher mutation frequency than the other two replicates (Figure S2). This suggests that the mutated *ftsI* contributed to the accelerated development of Amp-resistance under the co-exposure condition, and perhaps resulted in the higher Amp-resistance than the resistant mutants from Amp-exposure that developed different genetic mutations and resistance mechanisms.

### Differential Gene Expression of Resistant Mutants Isolated from Amp-exposed and Co-exposed *E. coli* Cultures

To further investigate the resistance mechanisms developed under co- and Amp-exposure conditions, differential gene expression analysis at the transcriptional level was conducted using RNA-seq. Principal component analysis indicates a clear difference between resistant mutants from co-exposure and those from Amp-exposure (Figure 4 A and B). A total of 92 and 107 genes exhibited significantly higher/lower expression (FDR < 0.05, ≥ 2-fold change) in Amp-R mutants and Cip-R mutants from co-exposure, respectively, compared to those from Amp-exposure. Hierarchical clustering revealed six distinct clusters of the differentially expressed genes under 8 functional categories. (Figure 4 C, D and details in Table S4).

**Fig. 4.**
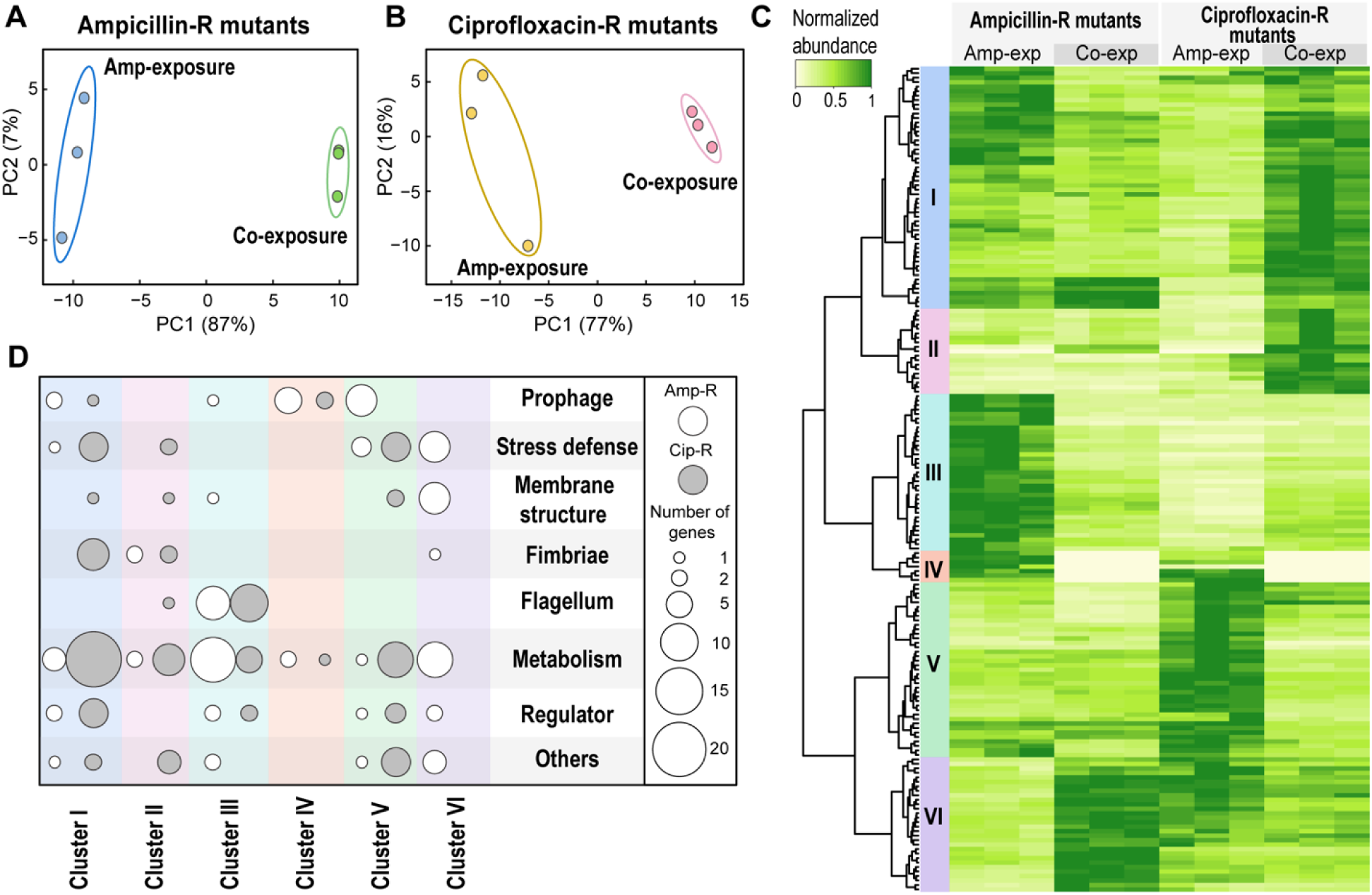
Differential gene expression analysis results, including principle component analysis of gene transcripts in Amp-R (A) and Cip-R (B) mutants from Amp-exposure and co-exposure after DESeq2 normalization, the heatmap of the relative abundance of differentially expressed genes (C), and the bubble plot of the number of differentially expressed genes in terms of gene clusters and gene functions (D).

Some genes in cluster I and almost all genes in cluster III showed significantly lower expression in Amp-R mutants from co-exposure than Amp-exposure, such as genes involved in flagellar structure formation (e.g., *fliC*), arginine synthesis (e.g., *argA*), carbohydrate transport (e.g., *argA, mglA*), cold shock defense (e.g., *cspH*), prophage (e.g., *nmpC, yjhQ*), and fatty acid β-oxidation (e.g., *fadB, fadH*). Moreover, the expression of genes in cluster IV was completely shut down in Amp-R mutants from co-exposure, including CP4-6 prophage genes (e.g., *yagE*, the mutated *yagJ*, and *mmuM* (also involving methionine synthesis)) and arginine synthesis genes (e.g., *argF*) (Figure S4A). In contrast, genes in cluster VI showed higher expression in Amp-R mutants from co-exposure, including heat shock and acid stress defense genes, such as *ibpA* and *hdeA*; genes involved in glutamate decarboxylation (*gadA, gadB*, and *gadC*), putrescine degradation (e.g., *puuB*) and histidine synthesis (*hisA* and *hisF*); and a membrane structure gene (*yhiM*) (Figure S4A). Two fimbriae-associated genes, *fimB* and *fimE*, in cluster II also showed higher expression levels in Amp-R mutants from co-exposure.

The Cip-R mutants from co-exposure exhibited higher expression of most genes in cluster I and II (Figure 4 C and Figure S4 B), including genes related to polymycin resistance (*pmrD* and *arnF*), heat shock defense (*patZ* and *ygcP*), oxidative stress defense (*bsmA*), histidine synthesis (e.g., *hisJ* and *hisM*), glyoxylate cycle (e.g., *aceA*), and N-acetylneuraminate degradation (e.g., *nanA*). In contrast, the expression of genes in cluster V, such as those associated with nitrate reduction (e.g., *narV*) and lipid degradation (*pagP* and *hdhA*) (Figure 4C and Figure S4B), was substantially lower in Cip-R mutants from co-exposed culture. The above gene expression patterns were quite different from those for Amp-R mutants, suggesting different resistance mechanisms. Interestingly, exceptions are found for two genes (i.e., *fimB* and *fimE*) encoding fimbrial structures, whose expression was stimulated in mutants resistant to both Amp and Cip from co-exposure. Besides, genes in cluster IV (e.g., prophage genes *yagJ, mmuM*) exhibited similar expression in both Amp-R and Cip-R mutants, which were turned off in mutants from co-exposed *E. coli* (Figure 4 C). These shared responses of the mutants from the co-exposure condition suggest the involvement of those genes in the resistance both to Amp and Cip. In addition, among all mutated genes identified in Amp-R and Cip-R mutants, *yagJ* was the only one that exhibited a significantly differential expression, in which there were several shared site mutations between the Amp- and Cip-R mutants from the co-exposure condition.

## Discussion

This work provides evidence that long-term exposure to pesticides alone or together with sub-MIC level antibiotics can stimulate and diversify *de novo* mutations towards resistance of certain antibiotics. The findings are of high relevance to the emergence of antibiotic resistance in some natural and built environments. High pesticide levels (mg/L) triggering evolution towards resistance may occur in biosolids and aquatic organisms where pesticides can be accumulated (49-51). In aquatic environments receiving WWTP effluent and agricultural runoff, antibiotics at sub-MIC levels are occurring together with pesticides at ng - µg/L (20-24). Such co-occurrence may synergistically select for *de novo* mutants resistant to antibiotics from a susceptible population, with even higher resistance than those that could have been selected by antibiotic exposure alone.

Mutation in genes encoding antibiotic target proteins is one of the mechanisms leading to higher resistance of mutants from pesticide-exposed and co-exposed *E. coli.* The higher resistance to Strep for mutants from pesticide-exposure was attributed to the stop-gain mutation in *rpsG* at a later amino acid position. The *rpsG* gene encodes a component (protein S7) of the 30S subunit of ribosome, and Strep binds to the 30S subunit to inhibit protein synthesis. This is different from previous findings that several site mutations in *rpsL*, another gene in the same operon encoding 30S subunit ribosomal protein S12, can lead to the structure alteration of 30S subunit, thus Strep-resistance in *E. coli* strains (52-54). The genetic change of *rpsG* uniquely selected under pesticide exposure may alter the structure of S7 and the entire 30S subunit, resulting in lower affinity, hence less sensitivity to Strep in the *de novo* mutants. Mutations in the antibiotic target genes, *ftsI* and *gyrA* for Amp and Cip, respectively, occurred exclusively (for *ftsI*) or more frequently (for *gyrA*) in resistant mutants from co-exposure than those from Amp-exposure. Direct alteration of the target proteins can be more effective to overcome the inhibitory effect of antibiotics than mutations in other resistance-related genes in the resistant mutants from Amp-exposure, leading to higher antibiotic resistance (MIC levels).

Moreover, the co-exposure to pesticides and Amp stimulated and diversified genome-wide mutations, and mutants with diverse mutations were selected under Cip stress, thus likely contributing to the higher Cip-resistance. Common mutations in a prophage gene *yagJ* were shared in both Amp- and Cip-resistant mutants from co-exposure, but not from Amp-exposure. The mutated gene *yagJ* exhibited differential expression (i.e., a complete shutdown) in both Amp- and Cip-resistant mutants from co-exposure compared to Amp-exposure. This differs from the previous findings that the removal of prophage CP4-6 genes including *yagJ* decreased the resistance to nalidixic acid (55), which is a quinolone antibiotic, as Cip is.

Previous studies (14, 15, 47) about the resistance mechanisms mostly focused on genetic mutations and the expression of antibiotic resistance genes. The global differential gene expression has not been well understood. Compared to the resistant mutants from Amp-exposure grown with antibiotic stress, the resistant mutants from co-exposure showed differential expression of many genes involved in metabolic activities and cell structure formation. Such different transcriptional responses to the same antibiotic stress may be related to the higher antibiotic resistance observed for the mutants from co-exposure than those from Amp-exposure. Amp- and Cip-R mutants from co-exposure shared several gene expression patterns, including (i) the stimulated expression of fimbriae synthesis genes promoting cell adhesion, and (ii) the deactivated expression of CP4-6 prophage-related genes, including *yagJ, ykgS*, and *mmuM*. These features may promote bacterial survival under stress conditions, rendering multidrug resistance.

In addition, we validated the differential gene expression results by RNA-seq using RT-qPCR targeting selected genes (Figure S5). According to RT-qPCR results, we also found that the differential expression of some genes in resistant mutants from co-exposure was independent of whether they were grown with antibiotic stress or not. For example, *fimB* and *fliC* in resistant mutants from co-exposure showed higher expression levels compared to the resistant mutants from Amp-exposure even when growing without antibiotic stress (Figure S6). This suggests that the distinct genetic mutations found in resistant mutants from co-exposure directly led to some transcriptional regulation without an antibiotic stimulus.

Taken together, this study unravels an overlooked role of pesticides in promoting the emergence of resistance to some antibiotics and selecting more resistant mutants with and/or without the presence sub-MIC level antibiotics. It gives a better understanding of the molecular mechanisms leading to the higher antibiotic resistance in *E. coli* after being exposed to multiple selection pressures rather than to antibiotics alone. This provides important insights into antibiotic resistance developed under more environmentally relevant exposure conditions.

## Materials and Methods

### Bacterial Strains, Growth and Selection Conditions

The antibiotic susceptible bacterium used in this study was the gram-negative *Escherichia coli* K-12 C3000 (*E. coli*). The growth medium for all selection experiments was Luria-Bertani (LB) broth. First, the stock *E. coli* cells from -80 °C freezer were revived and then streaked on a n LB agar plate and allowed to grow for 20 hours. One single colony was picked and inoculated into a tube containing 3 mL of LB broth for 24-hour incubation at 30 °C. The cell culture was considered as the ancestor strain and used for subsequent exposure experiments.

Twenty-three pesticides that have been frequently detected in environmental samples were selected. Their environmental concentrations (EC) range from 0.1 to 4.8 μg/L. Detailed information of the selected pesticides is in Table S1. Two exposure experiments were conducted: (1) Exposure to pesticide mixture of 1/125EC, 1/25EC, 1/5EC, 1EC, 5EC, 25EC, 125EC, mimicking a wide range of the pesticide occurrence in various environments with degradation or accumulation of pesticides. A no chemical exposure was also set up as the control. (2) Exposure to a combination of ampicillin of 1/125MIC_0_, 1/25MIC_0_, or 1/5MIC_0_ (MIC_0,_ MIC of antibiotics for the G0 *E. coli* strain in LB medium, MIC_0, Amp_ = 4 mg/L) and pesticides of 1EC was applied. The corresponding control was exposure only to ampicillin.

The pesticide stock mixture was dissolved in methanol. Appropriate volumes of the mixture were added to the 96-well plate, which was air-dried until all the methanol was gone. 195 μL LB medium and 5 μL ampicillin stock solution were subsequently added to the wells. The negative control group was added with the same volume of nanopure water as the ampicillin solution. The cultures were incubated at 30 °C and aerated by shaking. The culture was serially passaged by 500-fold dilution every 24 hours for 500 generations (9 generations of growth per serial passage). All exposure conditions and controls were performed with triplications.

### Isolation of Resistant Strains and Determination of Minimum Inhibitory Concentrations

The MIC_0_ of the ancestor strain was determined by the MIC test applied to 5 different types of antibiotics: ampicillin, tetracycline, ciprofloxacin, streptomycin, and chloramphenicol. Briefly, 95 μL of LB medium and 5 μL of the antibiotic stock solution were added, respectively. An overnight culture was prepared and diluted with 0.9% NaCl solution to optical density at 600 nm (OD_600_) of around 0.1 as the standard solution. Then 0.5 μL of the standard solution was added into fresh LB medium containing antibiotics at a series of concentrations. For the growth control group, 5 μL of nanopore water was used instead of the antibiotic solution. For the negative control group, 5 μL of nanopore water was used instead of the antibiotic solution and no *E. coli* was inoculated. Cell culture was incubated at 35 °C for 20 hours and OD_600_ were measured. The MIC was determined as the concentration that inhibited 90% of growth based on OD_600_.

After 500 generations, 5-time diluted cell cultures were spread on LB agar plates containing antibiotics at MIC_0_. Twelve resistant mutants were randomly picked up, and in total there were 36 resistant mutants from each exposure condition. The MICs of these resistant mutants were further determined. The Mann-Whitney U test was used to statistically analyze the difference of MICs among resistant mutants under different exposure conditions (*p*-value < 0.05).

### DNA Extraction, Whole-Genome Sequencing (WGS), and SNP Calling

Different *E. coli* isolates were cultured overnight in LB media and cell pellets were collected by centrifugation. DNA was extracted from each isolate using the DNeasy Blood and Tissue Kit (Qiagen) according to the manufacturer’s instructions. The DNA concentration and quality were determined on a Qubit 4 Fluorometer (Thermo Fisher Scientific, Wilmington, DE).

The obtained DNA obtained was then subjected to Illumina MiSeq 250-bp paired-end sequencing carried out by Roy J. Carver Biotechnology Center at the University of Illinois. An average coverage of 961,648 reads per isolate was obtained. A dynamic sequence trimming was done by SolexaQA software (56) with a minimum quality score of 24 and a minimum sequence length of 50 bp. The trimmed reads of the ancestor isolate (G0) were aligned against the *E. coli* K12 MG1655 genome available at NCBI GenBank (NC_000913.3) using the Bowtie 2 toolkit (57) to assemble the genome of *E. coli* at G0. All reads from isolates after G500 were then aligned against the assembled G0 genome. SAMtools and Picard Tools were used to format and reformat the intermediate-alignment files (58). SNPs and INDELs were identified with the Genome Analysis Toolkit UnifiedGenotyper (59), with the calling criteria of > 5-read coverage and > 50% mutation frequency at the mutation position.

### RNA Extraction, RNA-Seq, and Differential Gene Expression Analyses

One Amp-resistant mutant strain and one Cip-resistant mutant strain from different exposure conditions were selected from sequenced mutants. Mutants were grown in a shaking incubator at 35 °C in 8 mL LB broth for 5 hours to OD_600_ = 0.75. Each condition had 3 biological replicates. The cultures then were divided into two, one aliquot with the stress of 0.8×MIC antibiotic, one aliquot without antibiotic treatment. Cultures were allowed to grow for an additional 30 minutes, then cell pellets were collected by centrifugation.

Total RNA was isolated according to the acid phenol: chloroform extraction method, as previously described (60) and treated with DNase to remove residual DNA using TURBO DNA-free kit (Thermo Fisher Scientific). Ribosomal RNA was removed and sample libraries of resistant mutants with antibiotic treatment were built using a Truseq mRNA-Seq Library Preparation Kit (Illumina, USA), according to the manufacturer’s recommendations. Sequencing was performed on a HiSeq 2500 system (Illumina, USA) and produced 100-base single-end reads. The purified RNA samples of resistant mutants without antibiotic treatment, as well as those of G500 susceptible strains were reverse-transcribed to cDNA and stored properly for RT-qPCR measurement (See Supplementary Methods).

Low-quality RNA-seq reads (quality score < 30, sequence length < 25 bp) were removed using SolexaQA software (56). The qualified sequences were subject to the alignment using Bowtie 2 toolkit against the reference genome. Genes were counted using FeatureCounts software (61), and the count data were then analyzed using R version 3.5.1 and Bioconductor package DESeq2 version 3.8 (62). Genes were considered significantly differentially expressed based on these three criteria: (a) TPM (Transcripts per million) > 5 in at least one of the samples; (b) FDR (False Discovery Rate) adjusted *p*-value < 0.05; (c) > 2-fold difference in TPM values.

Principle component analysis was performed using normalized counts according to the DESeq2 output. Hierarchical clustering by transforming the normalized count data was applied based on the correlation distance and Ward aggregation criterion.

## Accession numbers

All WGS and RNA sequencing data have been deposited in the NCBI SRA database under accession no. PRJNA530028.

## Acknowledgements

We would like to give thanks to Hernandez Alvaro Gonzalo and Chris L. Wright at the Roy J. Carver Biotechnology Center, University of Illinois at Urbana-Champaign for whole genome sequencing and RNA sequencing support.

## References

1. Andersson DI & Hughes D (2014) Microbiological effects of sublethal levels of antibiotics. Nat Rev Microbiol 12(7):465–478.

2. Davies J & Davies D (2010) Origins and evolution of antibiotic resistance. Microbiol Mol Biol Rev 74(3):417–433.

3. Gullberg E, Cao S, Berg OG, Ilbäck C, Sandegren L, Hughes D, & Andersson D I (2011) Selection of resistant bacteria at very low antibiotic concentrations. PLoS Pathog 7:e1002158.

4. Long H, Miller SF, Strauss C, Zhao C, Cheng L, Ye Z, Griffin K, Te R, Lee H, Chen C-C, & Lynch M (2016) Antibiotic treatment enhances the genome-wide mutation rate of target cells. Proc Natl Acad Sci USA 113(18):E2498–2505.

5. Andersson DI & Hughes D (2011) Persistence of antibiotic resistance in bacterial populations. FEMS Microbiol Rev 35(5):901–911.

6. Brown KD, Kulis J, Thomson B, Chapman TH, & Mawhinney DB (2006) Occurrence of antibiotics in hospital, residential, and dairy effluent, municipal wastewater, and the Rio Grande in New Mexico. Sci Total Environ 366(2):772–783.

7. McArdell CS, Molnar E, Suter MJF, & Giger W (2003) Occurrence and fate of macrolide antibiotics in wastewater treatment plants and in the Glatt Valley watershed, Switzerland. Environ Sci Technol 37(24):5479–5486.

8. Watanabe N, Bergamaschi BA, Loftin KA, Meyer MT, & Harter T (2010) Use and environmental occurrence of antibiotics in freestall dairy farms with manured forage fields. Environ Sci Technol 44(17):6591–6600.

9. Li Y, Wu X, Mo C, Tai Y, Huang X, & Xiang L (2011) Investigation of sulfonamide, tetracycline, and quinolone antibiotics in vegetable farmland soil in the Pearl River Delta area, southern China. J Agric Food Chem 59(13):7268–7276.

10. Su J, Wei B, Ouyang W, Huang F, Zhao Y, Xu H, & Zhu Y (2015) Antibiotic resistome and its association with bacterial communities during sewage sludge composting. Environ Sci Technol 49(12):7356–7363.

11. Karkman A, Do TT, Walsh F, & Virta MPJ (2018) Antibiotic-resistance genes in waste water. Trends Microbiol 26(3):220–228.

12. Qian X, Sun W, Gu J, Wang X-J, Sun J-J, Yin Y-N, & Duan M-L (2016) Variable effects of oxytetracycline on antibiotic resistance gene abundance and the bacterial community during aerobic composting of cow manure. J Hazard Mater 315:61–69.

13. Zhu Y, Zhao Y, Li B, Huang C, Zhang S, Yu S, Chen Y, Zhang T, Gillings MR, & Su J (2017) Continental-scale pollution of estuaries with antibiotic resistance genes. Nat Microbiol 2:16270.

14. Lazar V, Nagy I, Spohn R, Csorgo B, Gyorkei A, Nyerges A, Horvath B, Voros A, Busa-Fekete R, Hrtyan M, Bogos B, Mehi O, Fekete G, Szappanos B, Kegl B, Papp B, & Pal C (2014) Genome-wide analysis captures the determinants of the antibiotic cross-resistance interaction network. Nat Commun 5:4352.

15. Kohanski MA, DePristo MA, & Collins JJ (2010) Sublethal antibiotic treatment leads to multidrug resistance via radical-induced mutagenesis. Mol Cell 37(3):311–320.

16. Lv L, Jiang T, Zhang S, & Yu X (2014) Exposure to mutagenic disinfection byproducts leads to increase of antibiotic resistance in *Pseudomonas aeruginosa*. Environ Sci Technol 48(14):8188–8195.

17. Li D, Zeng S, He M, & Gu AZ (2016) Water disinfection byproducts induce antibiotic resistance-role of environmental pollutants in resistance phenomena. Environ Sci Technol 50(6):3193–3201.

18. Baker-Austin C, Wright MS, Stepanauskas R, & McArthur JV (2006) Co-selection of antibiotic and metal resistance. Trends Microbiol 14(4):176–182.

19. Berg J, Thorsen MK, Holm PE, Jensen J, Nybroe O, & Brandt KK (2010) Cu exposure under field conditions coselects for antibiotic resistance as determined by a novel cultivation-independent bacterial community tolerance assay. Environ Sci Technol 44(22):8724–8728.

20. Kasprzyk-Hordern B, Dinsdale RM, & Guwy AJ (2009) The removal of pharmaceuticals, personal care products, endocrine disruptors and illicit drugs during wastewater treatment and its impact on the quality of receiving waters. Water Res. 43(2):363–380.

21. Kosma CI, Lambropoulou DA, & Albanis TA (2014) Investigation of PPCPs in wastewater treatment plants in Greece: occurrence, removal and environmental risk assessment. Sci Total Environ 466-467:421–438.

22. Luo Y, Guo W, Ngo HH, Nghiem LD, Hai FI, Zhang J, Liang S, & Wang XC (2014) A review on the occurrence of micropollutants in the aquatic environment and their fate and removal during wastewater treatment. Sci Total Environ 473-474:619–641.

23. Petrie B, Barden R, & Kasprzyk-Hordern B (2015) A review on emerging contaminants in wastewaters and the environment: current knowledge, understudied areas and recommendations for future monitoring. Water Res 72:3–27.

24. Bradley PM, Journey CA, Romanok KM, Barber LB, Buxton HT, Foreman WT, Furlong ET, Glassmeyer ST, Hladik ML, Iwanowicz LR, Jones DK, Kolpin DW, Kuivila KM, Loftin KA, Mills MA, Meyer MT, Orlando JL, Reilly TJ, Smalling KL, & Villeneuve DL (2017) Expanded target-chemical analysis reveals extensive mixed-organic-contaminant exposure in U.S. streams. Environ Sci Technol 51(9):4792–4802.

25. Harman-Fetcho JA, Hapeman CJ, McConnell LL, Potter TL, Rice CP, Sadeghi AM, Smith RD, Bialek K, Sefton KA, Schaffer BA, & Curry R (2005) Pesticide occurrence in selected South Florida canals and Biscayne Bay during high agricultural activity. J Agric Food Chem 53(15):6040–6048.

26. Bortoluzzi EC, Rheinheimer DS, Gonçalves CS, Pellegrini JBR, Maroneze AM, Kurz MHS, Bacar NM, & Zanella R (2007) Investigation of the occurrence of pesticide residues in rural wells and surface water following application to tobacco. Química Nova 30:1872–1876.

27. Ccanccapa A, Masia A, Navarro-Ortega A, Pico Y, & Barcelo D (2016) Pesticides in the Ebro River basin: occurrence and risk assessment. Environ Pollut 211:414–424.

28. Robles-Molina J, Gilbert-López B, García -Reyes JF, & Molina-Díaz A (2014) Monitoring of selected priority and emerging contaminants in the Guadalquivir River and other related surface waters in the province of Jaén, South East Spain. Sci Total Environ 479:247–257.

29. Wei R, Ge F, Huang S, Chen M, & Wang R (2011) Occurrence of veterinary antibiotics in animal wastewater and surface water around farms in Jiangsu Province, China. Chemosphere 82(10):1408–1414.

30. McManus PS, Stockwell VO, Sundin GW, & Jones AL (2002) Antibiotic use in plant agriculture. Annu Rev Phytopathol 40(1):443–465.

31. McKenna M (2019) Antibiotics set to flood Florida’s troubled orange orchards. Nature 567(7748):302.

32. Heeb F, Singer H, Pernet-Coudrier B, Qi W, Liu H, Longrée P, Müller B, & Berg M (2012) Organic micropollutants in rivers downstream of the megacity Beijing: sources and mass fluxes in a large-scale wastewater irrigation system. Environ Sci Technol 46(16):8680–8688.

33. Calderón -Preciado D, Matamoros V, & Bayona JM (2011) Occurrence and potential crop uptake of emerging contaminants and related compounds in an agricultural irrigation network. Sci Total Environ 412-413:14–19.

34. Fairbairn DJ, Karpuzcu ME, Arnold WA, Barber BL, Kaufenberg EF, Koskinen WC, Novak PJ, Rice PJ, & Swackhamer DL (2016) Sources and transport of contaminants of emerging concern: a two-year study of occurrence and spatiotemporal variation in a mixed land use watershed. Sci Total Environ 551:605–613.

35. Schwarzenbach RP, Escher BI, Fenner K, Hofstetter TB, Johnson CA, von Gunten U, & Wehrli B (2006) The challenge of micropollutants in aquatic systems. Science 313(5790):1072–1077.

36. Xing Y, Yu Y, & Men Y (2018) Emerging investigators series: occurrence and fate of emerging organic contaminants in wastewater treatment plants with an enhanced nitrification step. Environ Sci Water Res Technol 4(10):1412–1426.

37. Chang D-E, Smalley DJ, & Conway T (2002) Gene expression profiling of *Escherichia coli* growth transitions: an expanded stringent response model. Mol Microbiol 45(2):289–306.

38. Denyer SP (1990) Mechanisms of action of biocides. Int Biodeterior 26(2):89–100.

39. Heath RJ, Rubin JR, Holland DR, Zhang E, Snow ME, & Rock CO (1999) Mechanism of triclosan inhibition of bacterial fatty acid synthesis. J Biol Chem 274(16):11110–11114.

40. Newton BA (1956) The properties and mode of action of the polymyxins. Bacteriol Rev 20(1):14–27.

41. Attmannspacher U, Scharf BE, & Harshey RM (2008) FliL is essential for swarming: motor rotation in absence of FliL fractures the flagellar rod in swarmer cells of *Salmonella enterica*. Mol Microbiol 68(2):328–341.

42. Miryala SK & Ramaiah S (2018) Exploring the multi-drug resistance in *Escherichia coli* O157:H7 by gene interaction network: a systems biology approach. Genomics:pii: S0888–7543.

43. Ma D, Alberti M, Lynch C, Nikaido H, & Hearst JE (1996) The local repressor AcrR plays a modulating role in the regulation of *acrAB* genes of *Escherichia coli* by global stress signals. Mol Microbiol 19(1):101–112.

44. Okusu H, Ma D, & Nikaido H (1996) AcrAB efflux pump plays a major role in the antibiotic resistance phenotype of *Escherichia coli* multiple-antibiotic-resistance (Mar) mutants. J Bacteriol 178(1):306–308.

45. Parkhill J, Dougan G, James KD, Thomson NR, Pickard D, Wain J, Churcher C, Mungall KL, Bentley SD, Holden MTG, Sebaihia M, Baker S, Basham D, Brooks K, Chillingworth T, Connerton P, Cronin A, Davis P, Davies RM, Dowd L, White N, Farrar J, Feltwell T, Hamlin N, Haque A, Hien TT, Holroyd S, Jagels K, Krogh A, Larsen TS, Leather S, Moule S, Ó’Gaora P, Parry C, Quail M, Rutherford K, Simmonds M, Skelton J, Stevens K, Whitehead S, & Barrell BG (2001) Complete genome sequence of a multiple drug resistant *Salmonella enterica* serovar Typhi CT18. Nature 413:848.

46. Helling RB & Kukora JS (1971) Nalidixic acid-resistant mutants of *Escherichia coli* deficient in isocitrate dehydrogenase. J Bacteriol 105(3):1224–1226.

47. Oz T, Guvenek A, Yildiz S, Karaboga E, Tamer YT, Mumcuyan N, Ozan VB, Senturk GH, Cokol M, Yeh P, & Toprak E (2014) Strength of selection pressure is an important parameter contributing to the complexity of antibiotic resistance evolution. Mol Biol Evol 31(9):2387–2401.

48. Jaffé A, Chabbert YA, & Derlot E (1983) Selection and characterization of beta-lactam-resistant *Escherichia coli* K-12 mutants. Antimicrob Agents Chemother 23(4):622–625.

49. Chopra AK, Sharma MK, & Chamoli S (2011) Bioaccumulation of organochlorine pesticides in aquatic system—an overview. Environ Monit Assess 173(1):905–916.

50. Kannan K, Kajiwara N, Watanabe M, Nakata H, Thomas NJ, Stephenson M, Jessup DA, & Tanabe S (2004) Profiles of polychlorinated biphenyl congeners, organochlorine pesticides, and butyltins in southern sea otters and their prey. Environ Toxicol Chem 23(1):49–56.

51. Mohapatra DP, Cledon M, Brar SK, & Surampalli RY (2016) Application of wastewater and biosolids in soil: occurrence and fate of emerging contaminants. Water Air Soil Pollut 227(3):77.

52. Wistrand-Yuen E, Knopp M, Hjort K, Koskiniemi S, Berg OG, & Andersson DI (2018) Evolution of high-level resistance during low-level antibiotic exposure. Nat Commun 9(1):1599.

53. Ozaki M, Mizushima S, & Nomura M (1969) Identification and functional characterization of the protein controlled by the streptomycin-resistant locus in *E. coli*. Nature 222(5191):333–339.

54. Funatsu G & Wittmann HG (1972) Ribosomal proteins: XXXIII. Location of amino-acid replacements in protein S12 isolated from *Escherichia coli* mutants resistant to streptomycin. J Mol Biol 68(3):547–550.

55. Wang X, Kim Y, Ma Q, Hong SH, Pokusaeva K, Sturino JM, & Wood TK (2010) Cryptic prophages help bacteria cope with adverse environments. Nat Commun 1:147.

56. Cox MP, Peterson DA, & Biggs PJ (2010) SolexaQA: at-a-glance quality assessment of Illumina second-generation sequencing data. BMC Bioinformatics 11(1):485.

57. Langmead B & Salzberg SL (2012) Fast gapped-read alignment with Bowtie 2. Nat Methods 9:357.

58. Li H, Handsaker B, Wysoker A, Fennell T, Ruan J, Homer N, Marth G, Abecasis G, Durbin R, & 1000 Genome Project Data Processing Subgroup (2009) The sequence alignment/map format and SAMtools. Bioinformatics 25(16):2078–2079.

59. McKenna A, Hanna M, Banks E, Sivachenko A, Cibulskis K, Kernytsky A, Garimella K, Altshuler D, Gabriel S, Daly M, & DePristo MA (2010) The genome analysis toolkit: a mapreduce framework for analyzing next-generation DNA sequencing data. Genome Res 20(9):1297–1303.

60. Men Y, Yu K, Bælum J, Gao Y, Tremblay J, Prestat E, Stenuit B, Tringe SG, Jansson J, Zhang T, & Alvarez-Cohen L (2017) Metagenomic and metatranscriptomic analyses reveal the structure and dynamics of a dechlorinating community containing *Dehalococcoides mccartyi* and corrinoid-providing microorganisms under cobalamin-limited conditions. Appl Environ Microbiol 83(8):e03508–03516.

61. Liao Y, Smyth GK, & Shi W (2014) FeatureCounts: an efficient general purpose program for assigning sequence reads to genomic features. Bioinformatics 30(7):923–930.

62. Love MI, Huber W, & Anders S (2014) Moderated estimation of fold change and dispersion for RNA-seq data with DESeq2. Genome Biol 15(12):550.

