## Supplementary material for "Exposure to environmental level pesticides stimulates and diversifies evolution in *Escherichia coli* towards greater antibiotic resistance": SI

**This PDF file includes:**

SI Materials and Methods
Figs. S1 to S6
Tables S1 to S4
SI References

### SI Materials and Methods

**Determination of Mutation Frequencies.** The mutation frequency was measured according to a previous study (1). Briefly, 100 uL 1: 5 diluted (in 0.8% NaCl solution) overnight grown E. coli was spread on LB agar plates containing antibiotics. The total resistant mutant count per microliter equals to the number of colonies × 5 × $10^{-2}$. To count the total cell number, the same overnight culture was 1 : 5 × $10^{6}$ diluted and 100 μL of the diluted culture was spread on LB agar plates. The total bacteria count per microliter equals to the number of colonies × 5 × $10^{4}$. Three replicates were performed. The mutation frequency was then calculated by the following equation:

$$\text{Mutation frequency} = \frac{\text{Total resistant mutant count}}{\text{Total bacteria count}}$$

**SNP Genotyping Assay of *ftsI* Mutation.** Customized rhAmp SNP genotyping assay for *ftsI* mutation was designed and purchased (Integrated DNA Technology, USA). Co-exposed and Amp-exposed cultures of generation 500 were treated with 4 mg/L ampicillin ($MIC_0$), and then cell pellets were centrifuged and collected after 16-hour incubation. Genomic DNA (gDNA) samples was extracted from the cell pellets, and SNP genotyping assays of these samples were performed on the qPCR instrument according to the manufacturer's instruction. The positive controls of *ftsI* allele 1 (mutated *ftsI*) and allele 2 (non-mutated *ftsI*) were gDNA extracted from the Amp-R mutant and the susceptible isolate, respectively whose genomic information has been obtained from the whole genome sequencing.

**Quantitative Reverse Transcription PCR.** A minimum of 2 μg of total RNA was subject to cDNA synthesis using Superscript III (Invitrogen). The reversely transcribed samples were then used to employ quantitative PCR (qPCR) with the SYBRgreen method. Primers used in our study were listed in Table S2. The amplification targets included mutated genes and differentially expressed genes from RNA sequencing data. Three biological replications of each sample were included and raw cycle threshold (Ct) values were averaged to obtain the mean Ct values. A relative quantification strategy was applied, with the utilization of reference gene, *rpoB*. In brief, delta Ct values (ΔCt) of target gene and reference gene in each sample was first calculated as follows: $\Delta Ct = Ct_{target} - Ct_{reference}$. The delta delta Ct values (ΔΔCt) was then calculated by the equation: $\Delta\Delta Ct = \Delta Ct_{co\text{-}exposure} - \Delta Ct_{Amp\text{-}exposure}$. The relative transcription ratio (R) of mutants from co-exposure to mutants from Amp-exposure then was determined by the equation: $R = 2^{-\Delta\Delta Ct}$.

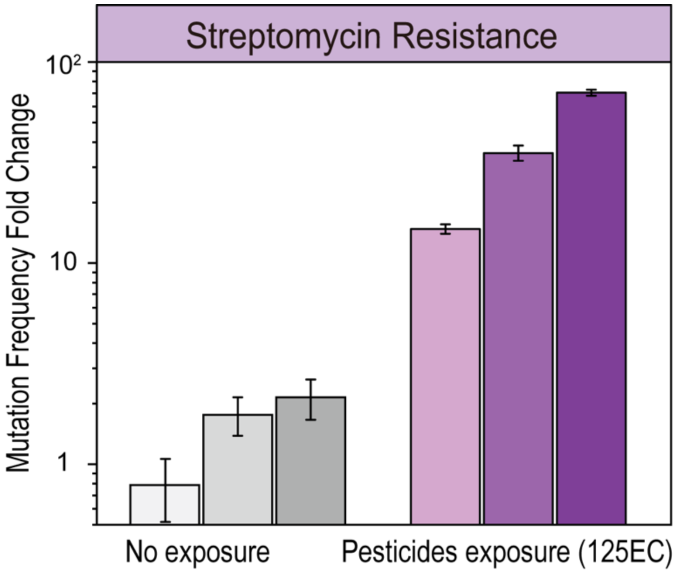


**Fig. S1.** Fold change of mutation frequency of *E. coli* towards streptomycin resistance under high-level pesticides-exposure and non-exposure conditions for 500 generations, comparing to that of the ancestor strain (G0). The mutation frequency was measured using three technical replicates for each of the three biological replicates, and the error bars represent the standard deviation of the three technical replicates.

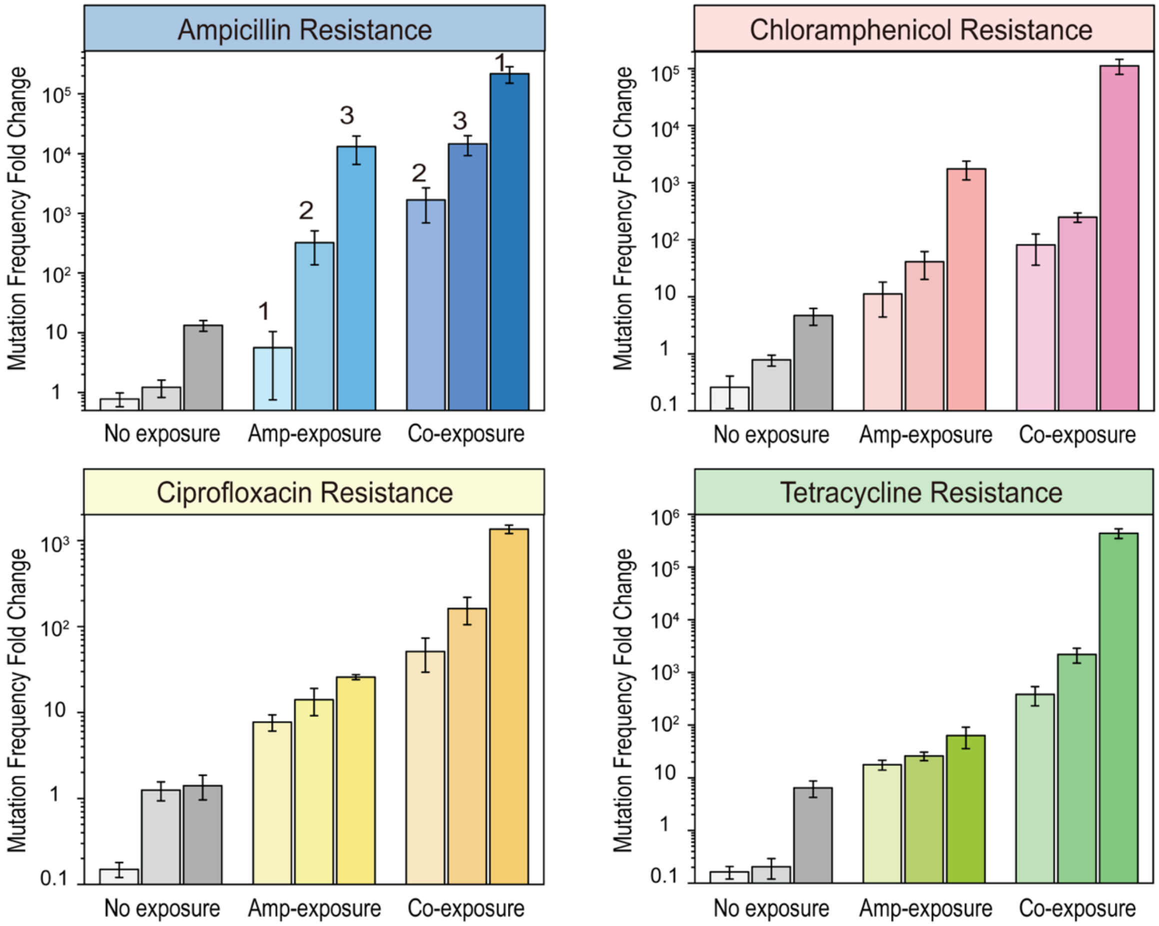


**Fig. S2.** Fold change of the mutation frequency of *E. coli* towards the resistance of ampicillin, chloramphenicol, ciprofloxacin and tetracycline, respectively under no exposure, single-exposure (Amp) and co-exposure (pesticides + Amp) for 500 generations (G500), comparing to the mutation frequency of the ancestor strain (G0). The mutation frequency was measured using three technical replicates for each of the three biological replicates. The columns represent averaged mutation frequencies of biological replicates, and the error bars represent the standard deviation of the three technical replicates for each biological replicate. The numbers of "1, 2, 3" in "Ampicillin Resistance" panel correspond to the same order of replicates in the SNP genotyping results (Fig. S3).

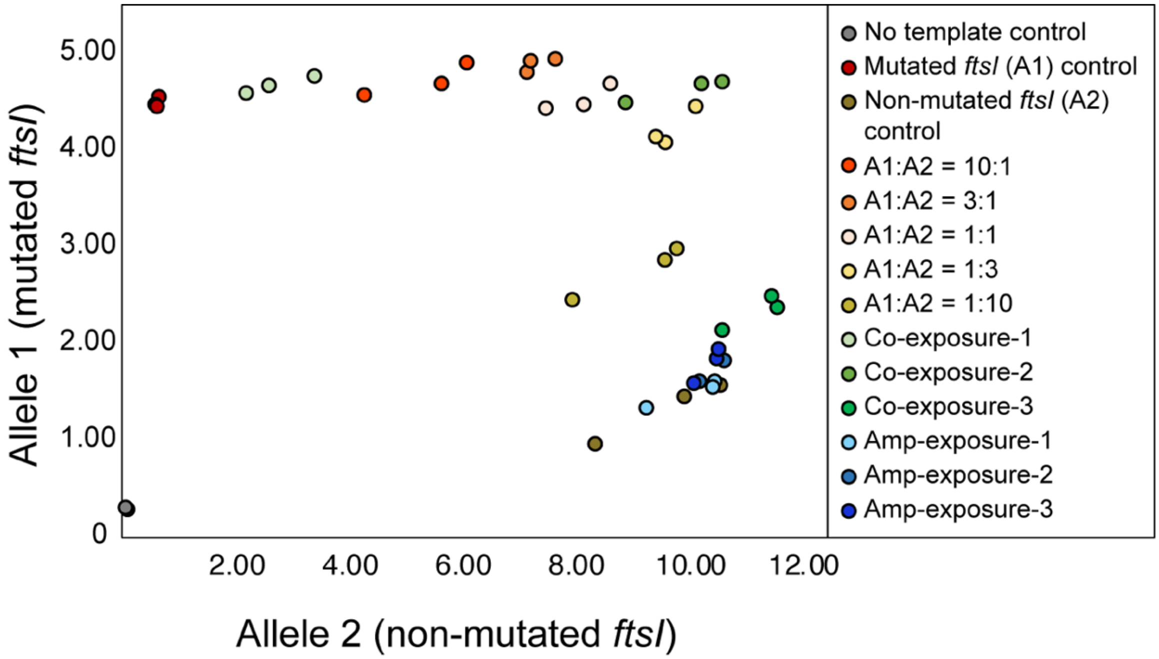


**Fig. S3.** Allelic discrimination plot of Amp-resistant cells from co-exposed (green circles) and Amp-exposed cultures (blue circles).

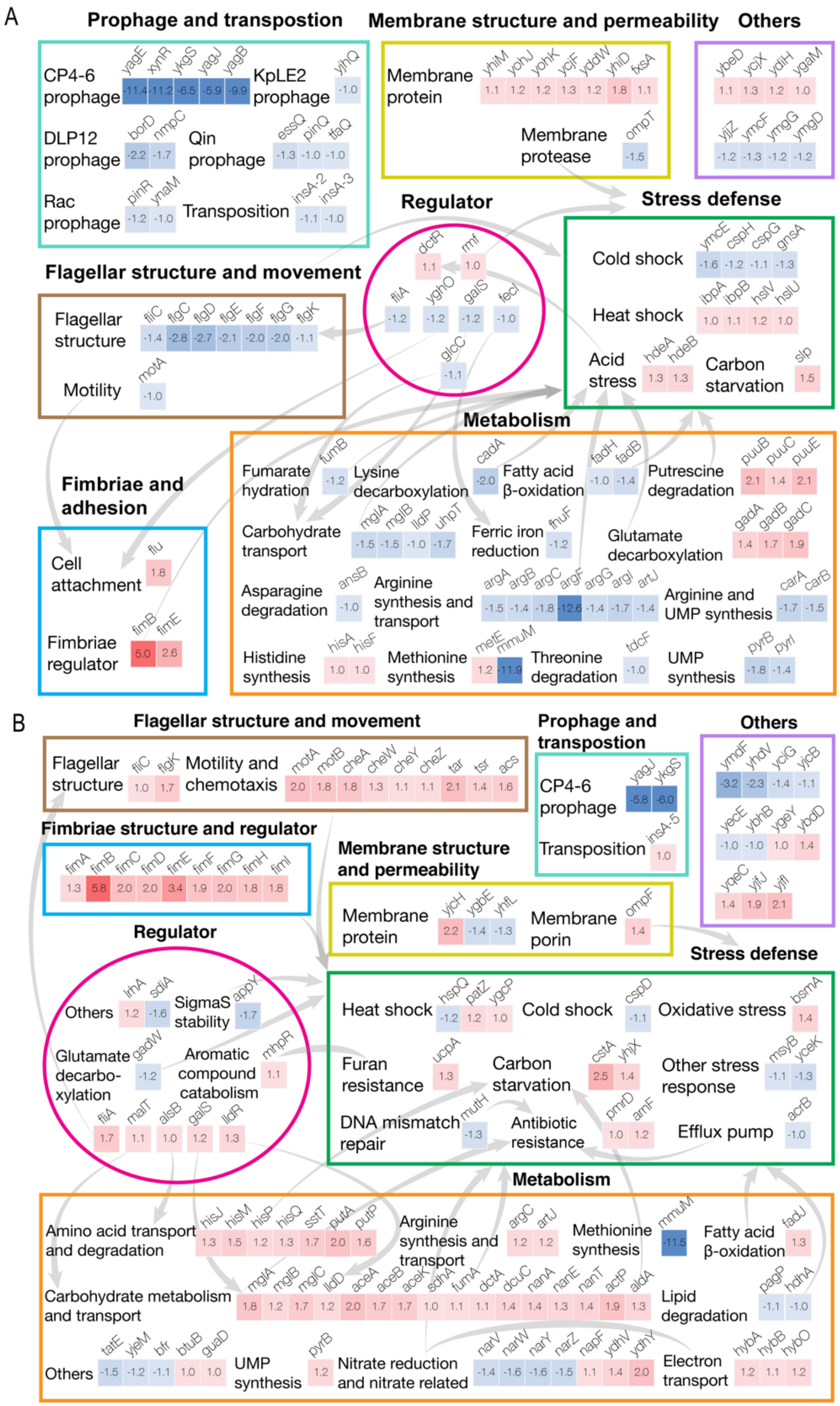


**Fig. S4.** Significantly differentially expressed genes in Amp-R mutants (A) and Cip-R (B) mutants selected from co-exposed *E. coli* in comparison with those from Amp-exposed *E. coli* grouped by gene functions. Numbers in each box indicate log2 difference of gene expression levels. Arrows link one gene to other gene(s) whose expression can be regulated by associated with the expression of the first gene according to literature (2-4).

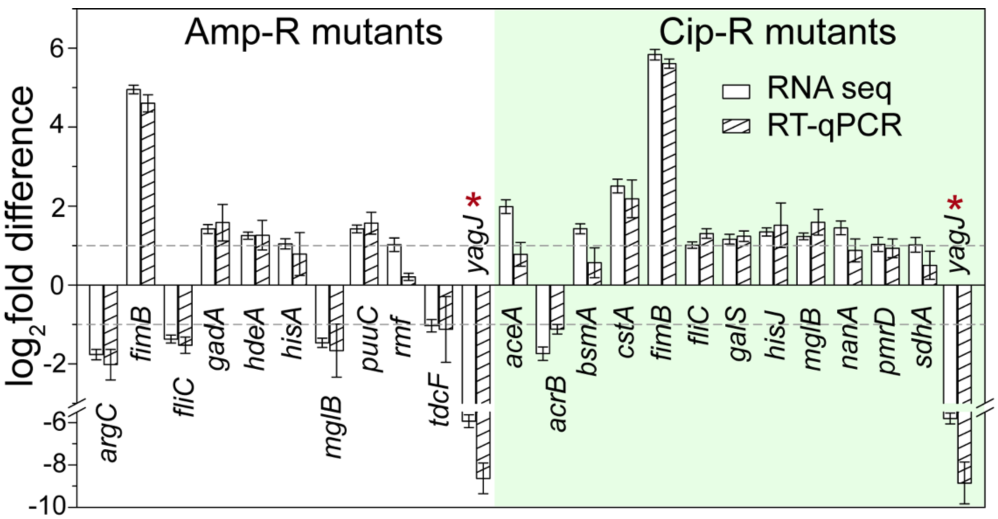


**Fig. S5.** Log2 fold difference between the expression of selected genes in resistant mutants from co-exposed *E. coli* and that from Amp-exposed *E. coli* with antibiotic stress comparing RNA-seq and RT-qPCR results. Genes with asterisk (*) indicates no expression (TPM < 5 for RNA-seq or Ct value > 30 for RT-qPCR) in resistant mutants from co-exposed *E. coli*.

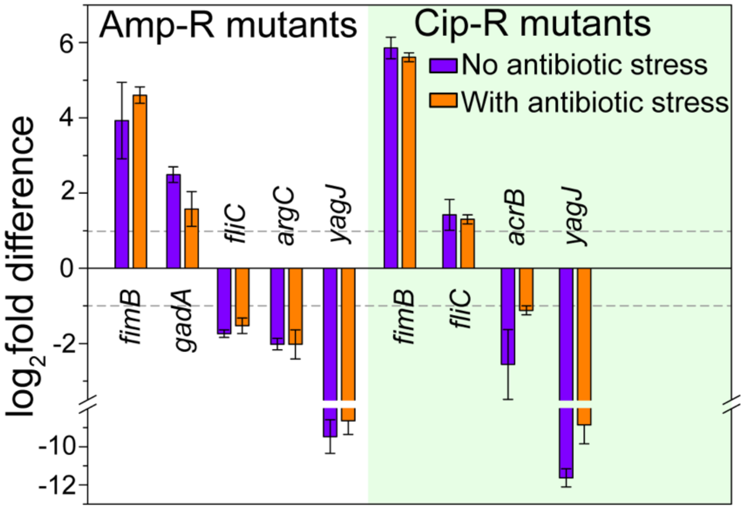


**Fig. S6.** Log2 fold difference between the expression of selected genes in resistant mutants from co-exposed *E. coli* and that from Amp-exposed *E. coli* without antibiotic stress (purple), and with antibiotic stress (orange).

**Table S1. Selected pesticides and their environmental concentrations (ECs).**

| Name | Classification | Conc. (μg/L) | Occurring environments | Standard Vender |
|---|---|---|---|---|
| 2,4-D | Herbicide | 0.2 | Urban run-off (5) | Sigma |
| Atrazine | Herbicide | 0.5 | Groundwater and surface water (6) | Sigma |
| Benomyl | Fungicide | 0.2 | Surface water (7) | Sigma |
| Carbaryl | Insecticide | 4.8 | Surface water (8) | Sigma |
| Carbofuran | Insecticide | 0.38 | Ground and surface water (9) | Sigma |
| Chlorpyrifos | Pesticide | 0.4 | Lake (10) | AK Scientific |
| Clotrimazole | Fungicide | 0.1 | Wastewater (11) | AK Scientific |
| DEET | Biocide | 3 | Wastewater influent (12) | Sigma |
| Diazinon | Insecticide | 0.3 | Wastewater (13) | Sigma |
| Diuron | Herbicide | 1 | Urban run-off (5) | AK Scientific |
| Fipronil | Insecticide | 0.2 | Urban surface water (14) | AK Scientific |
| Imazalil | Fungicide | 0.4 | River (15) | AK Scientific |
| Imidacloprid | Insecticide | 0.4 | Ground and surface water (9) | Sigma |
| Irgarol | Biocide | 0.2 | Coastal water (16) | Sigma |
| Linuron | Herbicide | 2 | Rivers (17) | Sigma |
| Mecoprop | Herbicide | 2 | Urban run-off (5) | Sigma |
| Metaldehyde | Pesticide | 0.5 | Surface water (18) | Sigma |
| Metolachlor | Herbicide | 0.4 | Wastewater (13) | Sigma |
| Propiconazole | Fungicide | 1 | Wastewater (19) | Sigma |
| Tebuconazole | Fungicide | 0.5 | Wastewater (20) | Sigma |
| Terbuthylazine | Herbicide | 0.65 | Ground and surface water (9) | AK Scientific |
| Terbutryn | Herbicide | 0.5 | Rivers (21) | Sigma |
| Thiabendazole | Fungicide | 0.2 | Wastewater influent (22) | Sigma |

**Table S2. Primers used in this study.**

| Target gene | Forward sequence (5' → 3') | Reverse sequence (5' → 3') |
|---|---|---|
| *aceA* | TTCCAGTTCATCACCCTGGC | GGCTGCTGCACTTTCTCAAC |
| *acrB* | CCCATCGACTTACGGGTAGC | ACAATGTTCGGGATGGTGCT |
| *argC* | GCAAAGCAATGATGCGGGAA | TGGCGAGAAACACTACGTCC |
| *bsmA* | TCGTCCACCATGACGACAAC | GACCAGACGCAAGGGTTACA |
| *cstA* | ATTGGCGAAGTGTCGGTCAT | GCCACAAAACCGTAACCCAC |
| *fimB* | GCCATTCGTGTGGTTTTGCT | CAGATGCCGTAAAAACGCCC |
| *fliC* | TCATCTCCGCCCAGTTTGAC | TACGCTGCGGATGTGAATGA |
| *gadA* | GGGCGAATTTATGCCAGCAG | ATGGCGATGAAATGGCGTTG |
| *galS* | TGGCCGATAATCCAGCTCAC | CATTCGTGATGTAGCGCGTC |
| *hdeA* | AAGCCTGAACGATAGCTGGG | CTGGCTGTGGACGAATCCTT |
| *hisA* | AAGAAGATGTGGCGGCGTTA | CACCGAAGCGTTCAAACCAG |
| *hisJ* | TCTTTAACAGACGGGCCACC | GACGTATTGATGCCGCGTTC |
| *mglB* | TTCTCAATCACCGTACCCGC | GACCAGTCCAAGCAGAACGA |
| *nanA* | GCAGACCAGAGGCGAAGATT | TACATTGCCTGGCGTAGGTG |
| *pmrD* | GCCACTTTACACTGCCGTTC | TGGGGGATTTACTCTCGCCT |
| *puuC* | TGTGGATGTGGACCCGAATG | AAGCTGTAGCGCCTGTTCTT |
| *rmf* | TCAACGTGGTTATCAGGCCG | CCATTACTACCCTGTCCGCC |
| *sdhA* | TTTCCCGTACCTTCGCTCAC | TGACCGGTAACTTTGGTCGG |
| *tdcF* | GTCGGATAGGTCGCCTGATG | CTGAGCGTGGGCGATATCAT |
| *yagJ* | AACCGCAGGATGTGGATCTG | CGGATAGTTCCCGCTTTCGT |
| *rpoB* | TCCGAAGGACAACCTGTTCG | AGGTCGAGGATCTGCTCTGT |

**Table S3. Complete genetic mutations identified in the resistant mutants.**

| Gene | Site position | Nucleotide change (SNP/INDEL) | Amino acid change | Samples | Gene function |
|---|---|---|---|---|---|
| ***rpsG*** | 3473665 | A → T | Stop gained (Leu157 → *) | Pesticide-exp, Strep-R-1, 2, 3 | 30S ribosomal subunit protein S7 |
| *arcB* | 3351726 | GA → G | Frameshift (Phe451 → fs) | Pesticide-exp, Strep-R-1 | Sensory histidine kinase |
| *sspA* | 3377241 | T → G | Tyr78 → Ser | Pesticide-exp, Strep-R-3 | Stringent starvation protein A |
| *rsmG* | 3923602 | ATCTCATTAGGAT → A | Non-frameshift deletion (Asp45_Met49 → Val) | Pesticide-exp, Strep-R-3 | 16S rRNA m(7)G527 methyltransferase |
| *proV* | 2805437 | A → T | Asp196 → Val | Amp-exp, Amp-R-1<br>Amp-exp, Cip-R-1 | Glycine betaine ABC transporter ATP binding subunit |
| *acrR* | 486033 | G → A | Glu91 → Lys | Amp-exp, Amp-R-2<br>Amp-exp, Cip-R-1 | DNA-binding transcriptional respressor |
| *icd* | 1196360 | C → CATCGAAAACATGTAATCTCTCCATGTGTTAAATAT | Frameshift and stop gained (Ala383 → fs) | Amp-exp, Amp-R-3 | Isocitrate dehydrogenase |
| ***ftsI*** | 93019 | A → T | Gln536 → Leu | Co-exp, Amp-R-1, 2, 3 | Peptidoglycan DD-transpeptidase |
| | 292171 | G → A | Val224 → Ile | Co-exp, Amp-R-1<br>Co-exp, Cip-R-3 | |

| | | | | | |
|---|---|---|---|---|---|
| ***yagJ*** | 292177 | A → C | Thr226 → Pro | Co-exp, Amp-R-1, 2, 3<br>Co-exp, Cip-R-2, 3 | CP4-6 prophage |
| | 292181 | C → A | Ala227 → Asp | Co-exp, Amp-R-1, 2, 3<br>Co-exp, Cip-R-2, 3 | |
| | 292186 | A → C | Asn229 → His | Co-exp, Amp-R-1, 2, 3<br>Co-exp, Cip-R-2, 3 | |
| | 292189 | G → GATCTCATAT | Disruptive inframe insertion (Ala230 → AspLeuIleSer) | Co-exp, Amp-R-1, 2, 3<br>Co-exp, Cip-R-2, 3 | |
| | 292192 | G → GGGACTTGTTC | Frameshift (Glu231 → fs) | Co-exp, Amp-R-1, 2, 3<br>Co-exp, Cip-R-2 | |
| | 292193 | A → G | Glu231 → Gly | Co-exp, Amp-R-1, 2, 3<br>Co-exp, Cip-R-2, 3 | |
| | 292194 | A → C | Glu231 → Asp | Co-exp, Amp-R-1, 2, 3<br>Co-exp, Cip-R-2, 3 | |
| | 292200 | A → C | Leu233 → Phe | Co-exp, Amp-R-1, 2, 3<br>Co-exp, Cip-R-2, 3 | |
| | 292201 | T → C | Phe234 → Leu | Co-exp, Amp-R-1, 2, 3<br>Co-exp, Cip-R-2, 3 | |
| *fliL* | 2019959 | C → A | Leu102 → Met | Co-exp, Amp-R-1 | Flagellar protein |
| *ydhJ* | 1721915 | A → G | Asp215 → Gly | Co-exp, Amp-R-3 | Putative membrane fusion protein |

| | | | | | |
|---|---|---|---|---|---|
| *yjcH* | 4285034 | A → G | Ile83 → Thr | Co-exp, Amp-R-3 | Conserved membrane protein |
| ***envZ*** | 3535564 | T → C | Thr120 → Ala | Amp-exp, Cip-R-1, 2 | Sensory histidine kinase |
| ***gyrA*** | 2339197 | T → C | Asp87 → Gly | Amp-exp, Cip-R-3<br>Co-exp, Cip-R-2 | DNA gyrase subunit A |
| | 2339209 | G → A | Ser83 → Leu | Co-exp, Cip-R-1, 3 | |
| *rluA* | 59991 | C → T | Stop gained<br>(Trp119 → *) | Co-exp, Cip-R-1 | 23S rRNA pseudouridine (746) and tRNA pseudouridine (32) synthase |
| *gsiA* | 868416 | GC → G | Frameshift<br>(Ile302 → fs) | Co-exp, Cip-R-1 | Glutathione ABC transporter ATP binding subunit |
| *aqpZ* | 915431 | A → G | Phe207 → Leu | Co-exp, Cip-R-1 | Water channel |
| *flgA* | 1130408 | A → G | Leu153 → Ser | Co-exp, Cip-R-1 | Flagellar basal body P-ring formation protein |
| *yneO* | 1594132 | GC → G | Frameshift<br>(Gly1321 → fs) | Co-exp, Cip-R-1 | AIDA-I family autotransporter Yneo |
| *fryA* | 2502562 | A → G | Cys654 → Arg | Co-exp, Cip-R-1 | Putative PTS multiphosphoryl transfer protein |
| *hyfB* | 2603798 | A → G | Ser649 → Gly | Co-exp, Cip-R-1 | Hydrogenase 4 component B |
| ***nlpD*** | 2867753 | A → AT | Frameshift<br>(Ile346 → fs) | Co-exp, Cip-R-1, 2, 3 | Murein hydrolase activator |

| | | | | | |
|---|---|---|---|---|---|
| *gudP* | 2921607 | A → G | Phe183 → Leu | Co-exp, Cip-R-1 | Galactarate transporter |
| *ebgC* | 3226222 | T → C | stop_lost<br>(Ter150 → Gln) | Co-exp, Cip-R-1 | DUF386 domain-containing evolved beta-D-galactosidase subunit beta |
| *ygjJ* | 3228130 | G → A | Gly93 → Ser | Co-exp, Cip-R-1 | Unknown |
| *tdcF* | 3259982 | A → G | Val61 → Ala | Co-exp, Cip-R-1 | Putative enamine/imine deasminase |
| *gpmM* | 3786242 | A → G | Tyr308 → Cys | Co-exp, Cip-R-1 | 2, 3-bisphosphoglycerate-independent phosphoglycerate mutase |
| ***epmB*** | 4375501 | A → G | Val76 → Ala | Co-exp, Cip-R-1, 2, 3 | Lysine 2, 3-aminomutase |
| *pmbA* | 4458500 | A → G | Thr158 → Ala | Co-exp, Cip-R-1 | Metalloprotease |
| *yjjP* | 4602348 | T → TG | Frameshift<br>(Thr194 → fs) | Co-exp, Cip-R-1 | Putative succinate exporter |
| *metQ* | 220867 | A → G | Val21 → Ala | Co-exp, Cip-R-2 | L-methionine/D-methionine ABC transporter membrane protein |
| *yafY* | 266380 | A → G | Val197 → Ala | Co-exp, Cip-R-2 | CP4-6 prophage |
| *mdlB* | 472339 | T → C | Tyr568 → His | Co-exp, Cip-R-2 | ABC transporter family protein |
| ***maa*** | 479493 | A → G | Val143 → Ala | Co-exp, Cip-R-2, 3 | Maltose O-acetyltransferase |
| ***sfmF*** | 563407 | A → G | Thr26 → Ala | Co-exp, Cip-R-2, 3 | Putative fimbrial protein |
| *cusS* | 593359 | T → C | Thr472 → Ala | Co-exp, Cip-R-2 | Sensory histidine kinase |

| | | | | | |
|---|---|---|---|---|---|
| ***ssuB*** | 993767 | A → G | Trp94 → Arg | Co-exp, Cip-R-2, 3 | Aliphatic sulfonate ABC transporter ATP binding subunit |
| ***dgcT*** | 1093354 | T → TG | Frameshift (Pro162 → fs) | Co-exp, Cip-R-2, 3 | Putative diguanylate cyclase |
| ***ydcI*** | 1495068 | AT → T | Frameshift (Asn4 → fs) | Co-exp, Cip-R-2, 3 | Putative DNA-binding transcriptional repressor |
| *adhP* | 1553732 | T → C | Thr39 → Ala | Co-exp, Cip-R-2 | Ethanol dehydrogenase |
| *yneK* | 1616197 | A → G | Thr143 → Ala | Co-exp, Cip-R-2 | Unknown |
| ***ynfM*** | 1670200 | A → G | Tyr165 → Cys | Co-exp, Cip-R-2, 3 | Putative transporter |
| *ydhV* | 1752004 | T → C | Tyr612 → Cys | Co-exp, Cip-R-2 | Putative oxidoreductase |
| *ynhG* | 1758328 | T → C | Thr136 → Ala | Co-exp, Cip-R-2 | L, D-transpeptidase |
| *ydiU* | 1790973 | G → A | Stop gained (Gln94 → *) | Co-exp, Cip-R-2 | UPF0061 family protein |
| ***btsS*** | 2214369 | GC → G | Frameshift (Gly104 → fs) | Co-exp, Cip-R-2, 3 | High-affinity pyruvate receptor |
| ***hcaD*** | 2672504 | T → C | Leu141 → Pro | Co-exp, Cip-R-2, 3 | Dioxygenase ferredoxin reductase subunit |
| *yqcE* | 2901090 | A → G | Thr155 → Ala | Co-exp, Cip-R-2 | Putative transport protein |
| *yqhG* | 3158138 | A → G | Ser146 → Gly | Co-exp, Cip-R-2 | DUF3828 domain-containing protein |
| *bfd* | 3467040 | T → C | Tyr2 → Cys | Co-exp, Cip-R-2 | Bacterioferritin-associated ferredoxin |

| | | | | | |
|---|---|---|---|---|---|
| *yhjJ* | 3681637 | T $\rightarrow$ G | Thr120 $\rightarrow$ Pro | Co-exp, Cip-R-2 | Peptidase M16 family protein |
| ***uvrA*** | 4273523 | A $\rightarrow$ G | Val139 $\rightarrow$ Ala | Co-exp, Cip-R-2, 3 | Excision nuclease subunit A |
| *ampC* | 4378143 | T $\rightarrow$ C | Asp291 $\rightarrow$ Gly | Co-exp, Cip-R-2 | Beta-lactamase |
| *djlB* | 678060 | A $\rightarrow$ G | Asp215 $\rightarrow$ Gly | Co-exp, Cip-R-3 | Putative chaperone |
| *nagA* | 702465 | G $\rightarrow$ A | Pro97 $\rightarrow$ Ser | Co-exp, Cip-R-3 | N-acetylglucosamine-6-phosphate deacetylase |
| *rutA* | 1073104 | T $\rightarrow$ C | Thr285 $\rightarrow$ Ala | Co-exp, Cip-R-3 | Pyrimidine oxygenase |
| *dhaR* | 1251903 | A $\rightarrow$ G | His279 $\rightarrow$ Arg | Co-exp, Cip-R-3 | DNA-binding transcriptional dual regulator |
| *adhE* | 1296185 | A $\rightarrow$ G | Cys647 $\rightarrow$ Arg | Co-exp, Cip-R-3 | Aldehyde-alcohol dehydrogenase |
| *trpE* | 1321762 | T $\rightarrow$ C | His398 $\rightarrow$ Arg | Co-exp, Cip-R-3 | Anthranilate synthase subunit |
| *ycjT* | 1379503 | A $\rightarrow$ G | Met538 $\rightarrow$ Val | Co-exp, Cip-R-3 | Kojibiose phosphorylase |
| *yhdP* | 3395758 | A $\rightarrow$ G | Val185 $\rightarrow$ Ala, no start codon | Co-exp, Cip-R-3 | AsmA2 domain-containing protein |
| *ubiD* | 4026208 | T $\rightarrow$ C | Val385 $\rightarrow$ Ala | Co-exp, Cip-R-3 | 3-octaprenyl-4-hydroxybenzoate decarboxylase |

**Table S4. Significantly differentially expressed genes in the 6 clusters in Figure 4.**

| Cluster 1 | | | Cluster 2 | | Cluster 3 | | | Class 4 | | Class 5 | | | Class 6 | |
|---|---|---|---|---|---|---|---|---|---|---|---|---|---|---|
| Amp-R | Cip-R | | Amp-R | Cip-R | Amp-R | | Cip-R | Amp-R | Cip-R | Amp-R | | Cip-R | Amp-R | |
| nmpC | insA-5 | fadJ | tdcF | acs | borD | motA | tsr | xynR | ykgS | ynaM | narW | ygbE | flu | hdeB |
| yjhQ | fimA | hisJ | ansB | yjcH | flgF | flgK | cheA | yagE | yagJ | pinR | bfr | yhfL | yhiD | hslV |
| fumB | fimC | hisM | fimE | aceA | flgE | artJ | cheW | yagB | mmuM | pinQ | appY | | yddW | hslU |
| fadH | fimD | hisP | fimB | aceB | flgG | pyrB | motB | argF | | essQ | sdiA | | yhiM | slp |
| fadB | fimF | hisQ | | aceK | flgD | mglA | tar | ykgS | | tfaQ | yceK | | fxsA | ygaM |
| lldP | fimG | hybB | | actP | flgC | mglB | fliC | yagJ | | insA-3 | msyB | | ycjF | ycjX |
| glcC | fimH | hybO | | putP | cheY | fliA | motA | mmuM | | insA-2 | cspD | | yohK | ybeD |
| yghO | fimI | lldD | | guaD | cheZ | galS | flgK | | | fhuF | mutH | | yohJ | ydiH |
| cspH | ompF | aldA | | ydhY | ompT | | artJ | | | fecI | acrB | | metE | |
| yjjZ | btuB | sstT | | ydhV | cadA | | pyrB | | | ymcE | hspQ | | puuB | |
| | napF | putA | | cstA | argB | | mglA | | | gnsA | ymdF | | puuC | |
| | hybA | mglC | | yhjX | argG | | mglB | | | cspG | ybhB | | puuE | |
| | dcuC | dctA | | ygeY | argC | | fliA | | | ymcF | yecE | | hisA | |
| | sdhA | nanA | | yjfJ | argA | | galS | | | gadW | yjcB | | hisF | |
| | mhpR | nanT | | yjfI | argI | | | | | yhdV | yciG | | gadA | |
| | patZ | nanE | | fimE | carA | | | | | narZ | | | gadB | |
| | pmrD | lldR | | fimB | carB | | | | | pagP | | | gadC | |
| | arnF | alsB | | | pyrI | | | | | tatE | | | dctR | |
| | ygcP | lrhA | | | uhpT | | | | | hdhA | | | rmf | |
| | ybdD | malT | | | ymgG | | | | | yjeM | | | ibpA | |
| | yqeC | ucpA | | | ymgD | | | | | narV | | | ibpB | |
| | fumA | bsmA | | | fliC | | | | | narY | | | hdeA | |

### References


1. Lv L, Jiang T, Zhang S, Yu X (2014) Exposure to mutagenic disinfection byproducts leads to increase of antibiotic resistance in *Pseudomonas aeruginosa*. *Environ Sci Technol* 48(14):8188-8195.
2. Franchini AG, Egli T (2006) Global gene expression in *Escherichia coli* K-12 during short-term and long-term adaptation to glucose-limited continuous culture conditions. *Microbiology* 152(7):2111-2127.
3. Chang D-E, Smalley DJ, Conway T (2002) Gene expression profiling of *Escherichia coli* growth transitions: an expanded stringent response model. *Mol Microbiol* 45(2):289-306.
4. Khil PP, Camerini-Otero RD (2002) Over 1000 genes are involved in the DNA damage response of *Escherichia coli*. *Mol Microbiol* 44(1):89-105.
5. Rippy MA*, et al.* (2017) Pesticide occurrence and spatio-temporal variability in urban run-off across Australia. *Water Res* 115:245-255.
6. Kolpin DW*, et al.* (2002) Pharmaceuticals, hormones, and other organic wastewater contaminants in U.S. streams, 1999−2000:  a national reconnaissance. *Environ Sci Technol* 36(6):1202-1211.
7. Senseman SA, Lavy TL, Mattice JD, Gbur EE, Skulman BW (1997) Trace level pesticide detections in Arkansas surface waters. *Environ Sci Technol* 31(2):395-401.
8. Elias D, Bernot MJ (2014) Effects of atrazine, metolachlor, carbaryl and chlorothalonil on benthic microbes and their nutrient dynamics. *PLoS One* 9(10):e109190.
9. Pitarch E*, et al.* (2016) Comprehensive monitoring of organic micro-pollutants in surface and groundwater in the surrounding of a solid-waste treatment plant of Castellón, Spain. *Sci Total Environ* 548-549:211-220.
10. Papadakis E-N*, et al.* (2015) Pesticides in the surface waters of Lake Vistonis Basin, Greece: occurrence and environmental risk assessment. *Sci Total Environ* 536:793-802.
11. Kahle M, Buerge IJ, Hauser A, Müller MD, Poiger T (2008) Azole fungicides: occurrence and fate in wastewater and surface waters. *Environ Sci Technol* 42(19):7193-7200.
12. Xing Y, Yu Y, Men Y (2018) Emerging investigators series: occurrence and fate of emerging organic contaminants in wastewater treatment plants with an enhanced nitrification step. *Environ Sci Water Res Technol* 4(10):1412-1426.
13. Campo J, Masiá A, Blasco C, Picó Y (2013) Occurrence and removal efficiency of pesticides in sewage treatment plants of four Mediterranean River Basins. *J Hazard Mater* 263:146-157.
14. Ensminger MP, Budd R, Kelley KC, Goh KS (2013) Pesticide occurrence and aquatic benchmark exceedances in urban surface waters and sediments in three urban areas of California, USA, 2008–2011. *Environ Monit Assess* 185(5):3697-3710.
15. Ccanccapa A, Masiá A, Navarro-Ortega A, Picó Y, Barceló D (2016) Pesticides in the Ebro River basin: occurrence and risk assessment. *Environ Pollut* 211:414-424.

16. Liu D, *et al.* (1999) Survey for the occurrence of the new antifouling compound Irgarol 1051 in the aquatic environment. *Water Res* 33(12):2833-2843.
17. Irace-Guigand S, Aaron JJ, Scribe P, Barcelo D (2004) A comparison of the environmental impact of pesticide multiresidues and their occurrence in river waters surveyed by liquid chromatography coupled in tandem with UV diode array detection and mass spectrometry. *Chemosphere* 55(7):973-981.
18. Hallett KC, Atfield A, Comber S, Hutchinson TH (2016) Developmental toxicity of metaldehyde in the embryos of *Lymnaea stagnalis* (Gastropoda: Pulmonata) co-exposed to the synergist piperonyl butoxide. *Sci Total Environ* 543:37-43.
19. Van De Steene JC, Stove CP, Lambert WE (2010) A field study on 8 pharmaceuticals and 1 pesticide in Belgium: removal rates in waste water treatment plants and occurrence in surface water. *Sci Total Environ* 408(16):3448-3453.
20. Stamatis N, Hela D, Konstantinou I (2010) Occurrence and removal of fungicides in municipal sewage treatment plant. *J Hazard Mater* 175(1):829-835.
21. Quednow K, Püttmann W (2007) Monitoring terbutryn pollution in small rivers of Hesse, Germany. *J Environ Monit* 9(12):1337-1343.
22. Guerra P, Kim M, Shah A, Alaee M, Smyth SA (2014) Occurrence and fate of antibiotic, analgesic/anti-inflammatory, and antifungal compounds in five wastewater treatment processes. *Sci Total Environ* 473-474:235-243.
